# Unraveling the functional role of DNA methylation using targeted DNA demethylation by steric blockage of DNA methyltransferase with CRISPR/dCas9

**DOI:** 10.1101/2020.03.28.012518

**Authors:** Daniel M. Sapozhnikov, Moshe Szyf

## Abstract

Although associations between DNA methylation and gene expression were established four decades ago, the causal role of DNA methylation in gene expression remains unresolved. Different strategies to address this question were developed; however, all are confounded and fail to disentangle cause and effect. We developed here a highly effective new method using only deltaCas9(dCas9):gRNA site-specific targeting to physically block DNA methylation at specific targets in the absence of a confounding flexibly-tethered enzymatic activity, enabling examination of the role of DNA methylation *per se* in living cells. We show that the extensive induction of gene expression achieved by TET/dCas9-based targeting vectors is confounded by DNA methylation-independent activities, inflating the role of DNA methylation in the promoter region. Using our new method, we show that in several inducible promoters, the main effect of DNA methylation is silencing basal promoter activity. Thus, the effect of demethylation of the promoter region in these genes is small, while induction of gene expression by different inducers is large and DNA methylation independent. In contrast, targeting demethylation to the pathologically silenced FMR1 gene targets robust induction of gene expression. We also found that standard CRISPR/Cas9 knockout generates a broad unmethylated region around the deletion, which might confound interpretation of CRISPR/Cas9 gene depletion studies. In summary, this new method could be used to reveal the true extent, nature, and diverse contribution to gene regulation of DNA methylation at different regions.

## Introduction

DNA methylation is broadly involved in transcriptional regulation across a vast number of physiological and pathological conditions [1]. For nearly half a century, it has been widely documented that the presence of methyl groups on the fifth carbon of cytosines in the context of CpG dinucleotides within promoters is associated with transcriptional repression [2]. This is considered to be a crucial epigenetic mark as deviations from the tightly-regulated and tissue-specific developmental patterns have been implicated in conditions as diverse as cancers [3], suicidal behavior [4], and autoimmune diseases [5]. Yet, these studies also exemplify a fundamental challenge in the field: the persistent inability to attribute causality to a particular instance of aberrant DNA methylation. The issue of whether DNA methylation is the driver of relevant transcriptional changes continues to be a source of controversy and is magnified by multiple studies suggesting that changes in gene expression and transcription factor binding can in some cases precede changes in DNA methylation [6–11]. The answers to this set of questions would reveal whether a particular DNA methylation state is only a marker for a particular condition or whether it is plays a critical role in the pathophysiological mechanism.

In the case of DNA methylation, unconfounded manipulation of the methylation state of a CpG or region of CpGs in isolation remains a challenge: genetic (DNA methyltransferase knockdown) and pharmacological (5-aza-2’-deoxycytidine and S-Adenosyl methionine) hypo- or hyper-methylating agents cause genome-wide changes in methylation[12–16], confounding conclusions by countless concurrent changes throughout the genome in addition to any region under study. A more specific approach to assessing causality involves comparing the abilities of *in vitro* methylated and unmethylated regulatory sequences to drive reporter gene expression in transient transfection assays. However, this is an artificial system and a simplification of the complex chromatin architecture at the endogenous locus, and therefore the effects of methylation in the context of an artificial promoter-reporter plasmid may misrepresent those that would occur under physiological conditions.

More recently, the TET dioxygenases – which oxidize the methyl moiety in cytosine and can lead to passive loss of methylation by either inhibiting methylation during replication or through repair of the oxidized methylcytosine and its replacement with an unmethylated cytosine – were targeted to specific sites using a fusion of TET dioxygenase domains to catalytically inactive CRISPR/Cas9 (deltaCas9 or dCas9) [17–20]. However, this method still introduces several confounding factors that preclude causational inferences, such as the fact that oxidized methylcytosines are new epigenetic modifications that are not unmethylated cytosines [21–27] and the fact that TET has methylation-independent transcriptional activation activity[28, 29]. We propose and optimize instead an enzyme-free CRISPR/dCas9-based system for targeted methylation editing which we show to be able to achieve selective methylation *in vitro* and passive demethylation in cells through steric interference with DNA methyltransferase activity. We map the size of the region of interference, optimize the system for nearly complete demethylation of targeted CpGs without detectable off-target effects, and analyze the transcriptional consequences of demethylation of genetically dissimilar regions across several human and mouse genes. In doing so, we provide evidence that DNA demethylation at proximal promoters increases gene expression in some instances but not others, that it does so to varying degrees depending on the genomic context, and that demethylation may facilitate responses to other factors. Most importantly, we report a simple tool for investigations into the effects of DNA methylation that can be applied with ease and in multiplexed formats to examine the vast existing and forthcoming correlational literature in order to distinguish causational instances of DNA methylation and begin to develop a fundamental understanding of this biological phenomenon on a foundation of causality.

## Results

### CRISPR/TET-based approaches confound the causal relationship of DNA methylation and transcription

To develop a tool for site-specific DNA methylation editing, we elected to study the murine interleukin-33 (*Il33*) gene. The distance between individual CpGs and sets of CpGs within its canonical CpG-poor promoter provides a simple starting point that enables specific CpG targeting in order to evaluate the impact of discrete methylation events on gene transcription (Fig. 1A). The promoter is highly methylated in NIH-3T3 cells and upon treatment of cells with the demethylating agent 5-aza-2’-deoxycytidine, CpGs adjacent to the transcription start site (TSS) are demethylated (Fig. 1B) and gene expression is moderately induced (Fig. 1C). However, this classical response to 5-aza-2’-deoxycytidine also emphasizes the shortcomings of this common approach in DNA methylation research: (1) multiple CpGs in the promoter are demethylated, so it remains unclear which sites of methylation contribute to transcriptional inhibition, and (2) the global genomic consequences of 5-aza-2’-deoxycytidine treatment result in the induction of expression of several putative and experimentally validated *Il33* transcription factors (Fig. 1D), exemplifying the possibility that demethylation of the *Il33* promoter may not be the event responsible for upregulation of the gene. This demonstrates a need for an accurate and specific targeted methylation editing technology that can properly interrogate the fundamental question of the causal relationship between DNA methylation at specific sites and gene expression in *cis*.

**Figure 1.**
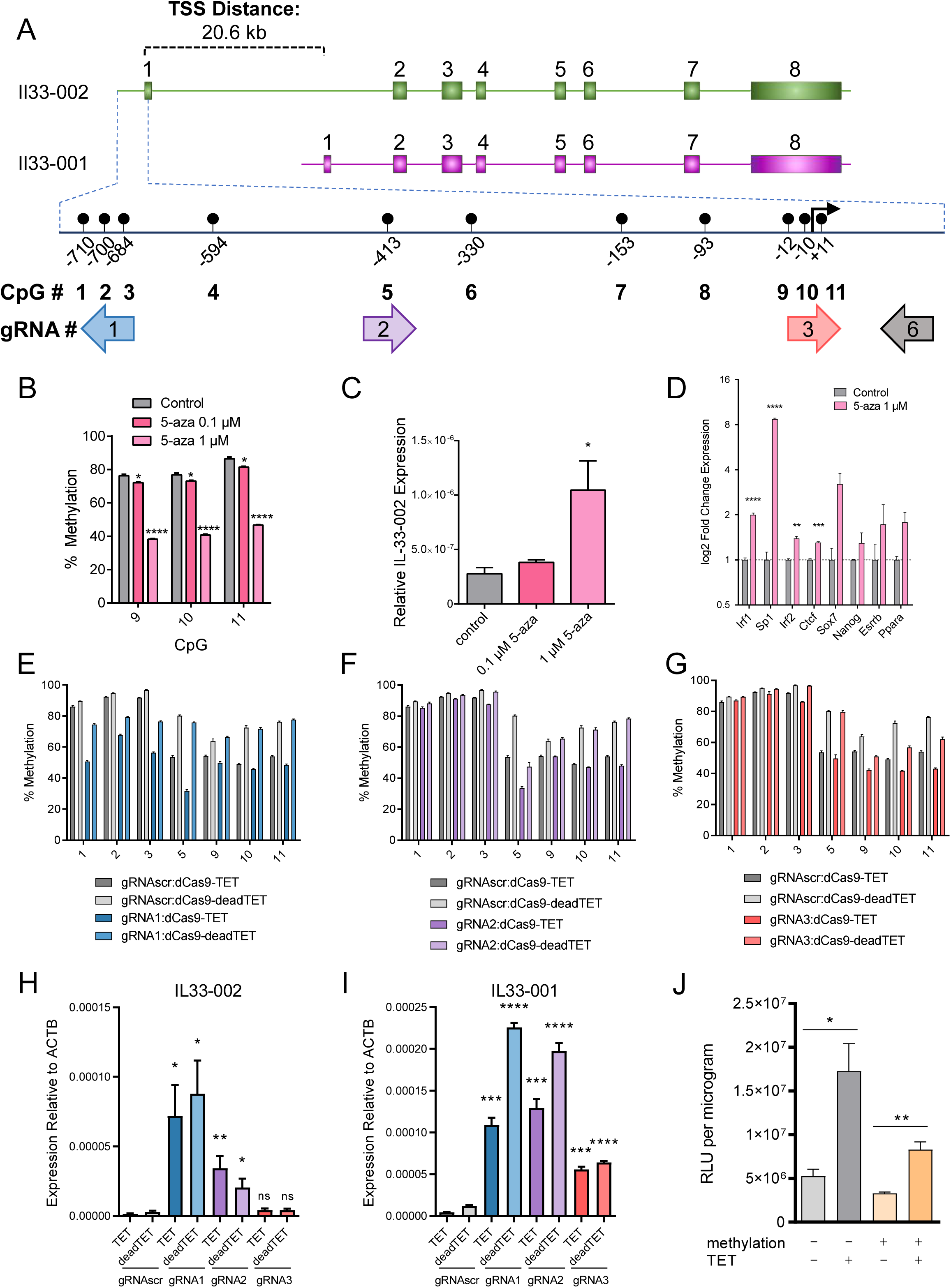
Targeting the Il33 promoter with dCas9-TET. (A) Schematic of the murine *Il33* genomic locus depicting the two transcriptional isoforms with a highlighted region of an 800bp region of the *Il33-002* promoter and the locations of the 11 CpGs as well as 4 gRNAs targeting specific CpGs. The 11 CpGs are numbered sequentially in the 5’ to 3’ direction. The promoter-targeting gRNAs used in these experiments are shown relative to the CpGs and are approximately to scale such that CpGs 1, 2, and 3 are targeted by gRNA1, CpG 5 by gRNA 2, and gRNA 3 targets CpGs 9, 10, 11 – which overlap the transcription start site (TSS), marked by a black arrow. The orientation of the gRNAs is indicated by an arrow, where an arrow pointing to the left indicates a gRNA that binds the plus strand. (B) Percent of DNA methylation assayed by bisulfite-pyrosequencing at the three transcription start site (TSS) CpGs (labeled 9-11) following treatment of NIH-3T3 cells with indicated concentrations of 5-aza-2’-deoxycytidine or water control. (C) Expression of *Il33-002* quantified by RT-qPCRand normalized to *beta Actin* (*Actb*) expression following treatment of NIH-3T3 cells with indicated concentrations of 5-aza-2’-deoxycytidine or water control. (D) Expression of predicted (Transfac) and experimentally validated (Qiagen, ENCODE, Gene Transcription Regulation Database) Il33-002 transcription factors quantified by RT-qPCR and normalized to *Actb* expression following treatment of NIH-3T3 cells with indicated concentrations of 5-aza-2’-deoxycytidine or water control. (E-G) Percent of DNA methylation assayed by bisulfite-pyrosequencing at 7 targeted CpGs in the *Il33-002* promoter following transduction with lentiviruses and antibiotic selection of virally infected cells (gRNAs) or selection by flow cytometry (BFP; dCas9 constructs) of NIH-3T3 cells with dCas9-Tet/dCas9-deadTET (BFP) and gRNA1 (E), gRNA2 (F), or gRNA3 (G) compared to gRNAscr (light and dark grey, data identical in E-G and shown for comparison) (n=4-8). (H-I) Expression of *Il33-002* (H) and *Il33-001* (I) quantified by RT-qPCR and normalized to *Actb* expression in NIH-3T3 stably expressing one of 4 gRNAs and dCas9-TET or dCas9-deadTET. n=3 for all data panels, except E-G as indicated, and all data shown as (mean+/-SEM). (J) Relative light units normalized to protein quantity in transfected HEK293 cells. Cells were transiently transfected with methylated or unmethylated SV40-luciferase vector along with mammalian TET2 expression plasmid or empty vector (pcDNA3.1) control. * indicates statistically significant difference of p<0.05, ** of p<0.01, *** of p<0.001, **** of p<0.0001, and ns = not significant (Student’s t-test, with Holm-Sidak correction if number of tests is greater than 3).

To first assess the efficacy and specificity of the available targeted DNA methylation editing technology, we examined the lentiviral system created by Liu et al [30] consisting of a catalytically inactive Cas9 (dCas9) fused to the catalytic domain of TET1 (dCas9-TET or a catalytic mutant, dCas9-deadTET), which is thought to promote active DNA demethylation by oxidation of the methyl moiety and eventual replacement of the modified cytosine with unmethylated cytosine by the base excision repair pathway [17]. We developed a set of 20 base-pair (bp) CRISPR guide RNAs (gRNAs) targeting distinct regions in the promoter of the *Il33-002* transcript, the inducible variant [31, 32] (Fig. 1A and Supp. Table S1). The system was effective in partially demethylating the *Il33* promoter; however, we noted several shortcomings of this method.

First, it was immediately apparent that NIH-3T3 cells expressing only a scrambled, non-targeting guide (gRNAscr) and dCas9-TET were significantly more demethylated than counterparts expressing the same non-targeting gRNA and dCas9-deadTET (Figure 1E-G), particularly at CpGs 5, 10, and 11 (range from ∼ 22 to 26 percent) but also to a smaller degree at all remaining evaluated CpGs. This is indicative of a potential ubiquitous and dCas9-independent activity of the fused, over-expressed TET domain. It stands to reason that overexpressed proteins would interact with several DNA targets across the genome by their cognate DNA binding domain in the absence of any additional targeting. Indeed, overexpression of DNMT3A [33, 34] and TET [35–37] without any targeting were previously shown to broadly alter DNA methylation. In fact, DNMT3A still deposits a genomic methylation pattern similar to DNMT3A overexpression alone despite specific targeting by dCas9 [38]. Given these data and our observations, we anticipate a similar ability of TET catalytic domains to interact with their cognate target sites despite flexible fusion to dCas9, inevitably leading to off-target effects that impede accurate conclusions about the causality of changes to DNA methylation.

Second, the demethylation caused by dCas9-TET spanned a substantial genetic distance. For example, in gRNA1:dCas9-TET cells, while the protein complex was positioned at and significantly demethylated CpGs 1, 2, and 3 (p<0.0001), the remaining CpGs were all significantly demethylated as well, including CpG 11 (p=0.00014), which is nearly 700 bp away from gRNA1. Similar significant long-distance demethylation effects could be observed in cells expressing gRNA2 and gRNA3. The potential for long-distance effects is further exemplified in the strong transactivation effects of dCas9-TET positioned at the *Il33-002* promoter on the distant *Il33-001* promoter, approximately 21 kb away (Fig. 1I). This illustrates a second category of confounding off-target effects that are introduced by a targeting strategy that employs a flexibly tethered enzyme which can modify genetic regions in close physical proximity – despite large genetic distances – particularly those in ubiquitous self-interacting topologically associating domains (TADs), such as the one that the two *Il33* promoters belong to [39].

Third, when evaluating the transcriptional effects of the epigenetic editing system, we were surprised to discover that dCas9-deadTET paired with gRNA 1 or 2 (gRNA3 blocks the TSS and likely interferes with RNA polymerase binding [40]) resulted in strong transactivation of the *Il33-002* transcript to levels comparable to dCas9-TET (Fig. 1H), despite lacking any catalytic capacity to initiate the active DNA demethylation process. These data appear to suggest that the TET1 domain possesses a transcriptional activation capacity that is independent of its DNA modification function, thereby confounding any inferences as to the causal relationship between DNA methylation *per se* and gene expression. Even more surprising was the fact that that dCas9-deadTET was more effective in transactivation at long distance (Fig. 1I) than dCas9-TET. The *Il33-001* transcript was significantly more expressed in dCas9-deadTET cells under gRNA1 (p=0.0091) and gRNA3 (p=0.0033) as compared to dCas9-TET. To ensure that this unexpected result was not a consequence of erroneous sample switches, we amplified the region containing the catalytic mutations of the TET1 domain in the DNA samples used for methylation analysis and in the cDNA samples used for expression quantification and confirmed by Sanger sequencing that all dCas9-deadTET samples bore the two point mutations that render it catalytically inactive (Supp. Fig. S1A).

To assess by a secondary measure the DNA methylation independent transactivation capacity of TET proteins, we performed transient co-transfections of *in vitro* methylated or unmethylated promoter-reporter plasmids – luciferase driven by the SV40 promoter/enhancer – in combination with a mammalian expression vector expressing the human TET2 catalytic domain (pcDNA-TET2) or an empty vector control. We found that the TET2 catalytic domain induces the activity of completely unmethylated promoters (Fig. 1J), reaffirming the notion that TET proteins produce transcriptional changes independently of any DNA demethylation and thus cofounding correlational assessments.

Additionally, we combined our three targeting gRNAs with the well-characterized dCas9-VP64 fusion; VP64 is a potent transcriptional activator originating from the herpes simplex virus [41]. The tetramer of the herpes simplex VP16 protein acts to activate transcription primarily through recruitment of basal transcription machinery, including TFIID/TFIIB, and has no known catalytic capacity for DNA demethylation [42]. Yet, we found that dCas9-VP64 co-expressed with all 3 *Il33-002* gRNAs resulted in dramatic and broad demethylation of the *Il33-002* promoter (Supp. Fig. S1B-D). This suggests that DNA demethylation can in particular instances be secondary to transcription factor recruitment and transcriptional activation (Supp. Fig. S1E.). We again found significant and robust activation of the distant *Il33-001* promoter (gRNA2, p<0.05; gRNA3, p<0.001), supporting the notion that enzymatic domains flexibly tethered to dCas9 can act across large genetic distances (Supp. Fig. S1F). Combined with the facts that dCas9-deadTET can strongly induce gene expression and the TET2 catalytic domain can induce unmethylated promoters, these data provide further evidence that the dCas9-TET strategy is highly confounded by the multifunctionality of the TET domain.

Finally, it is important to note that the TET proteins are not catalytic demethylases. They instead promote active demethylation through a pathway that contains several stable modified-cytosine intermediates (5-hydroxymethylcytosine, 5-formylcytosine, and 5-carboxylcytosine), all of which are uniquely recognized by different transcription factors and may participate in epigenetic recruitment with transcriptional consequences that confound conclusions concerning the causality of DNA demethylation events [21–23, 25, 43, 44]. We detected a significant increase of 5-hydroxymethylcytosine in the *Il33-002* promoter in the presence of dCas9-TET but not dCas9-deadTET (Supp. Fig. S1G), demonstrating that demethylation is not the only epigenetic change conferred by dCas9-TET.

Taken together, these data reveal that while dCas9-TET may be a valid tool for producing epigenetic perturbations that may further understanding of TET dynamics, it introduces a number of confounds inherent to the properties of the TET protein that prohibit conclusions as to the causal relationship of changes in DNA methylation at particular sites and gene expression.

### A novel method for site-specific DNA methylation *in vitro*

A potential mechanism for producing specific demethylation in cells is through targeted physical interference with the DNA methyltransferase (DNMT) machinery that deposits methyl groups onto nascent post-replicative DNA. We reasoned that since dCas9 is able to interfere with transcriptional machinery to reduce gene expression [40], it may also be able to sterically obstruct DNMT activity at its binding position. Since dCas9 is a prokaryotic protein that is unlikely to interact with other proteins in the gene transcription machinery and has no enzymatic activity epigenetic or other, there should be limited confounding factors to DNA demethylation.

To test this hypothesis, we first investigated whether dCas9 could be applied as a tool to interfere with DNMT activity at targeted CpGs in a simplified *in vitro* system. The target DNA used for methylation was a 1,015 bp fragment of the *Il33-002* promoter (Fig. 1A) inserted into an otherwise CpG-free luciferase reporter vector [45] to enable the assessment of methylation changes on reporter gene activity in transient transfection assays. Standard methylation with the bacterial CpG methyltransferase M.SssI protein resulted in 80-93% methylation at all CpGs as assayed by pyrosequencing (Fig. 2A) and a significant 4-fold decrease in luciferase reporter activity in a transient transfection assay (p=0.0041) (Fig. 2F). Incubation of *Il33*-pCpGl with recombinant dCas9 protein and an *in vitro* transcribed chimeric control gRNA (gRNA6 in Fig. 1A) targeting the CpG deficient region approximately 110-130bp downstream of the TSS only slightly inhibited the efficiency of the M.SssI reaction at all CpGs (Fig. 2A). The plasmid was still highly methylated and the treatment also significantly reduced luciferase activity (p=0.0018) compared to mock treatment and to a similar extent as standard methylation (p=0.374) (Fig. 2F). This confirms that the reaction components (including dCas9 protein, non-CpG-targeting gRNA, buffer system, and incubation times) do not compromise DNA methyltransferase activity.

**Figure 2.**
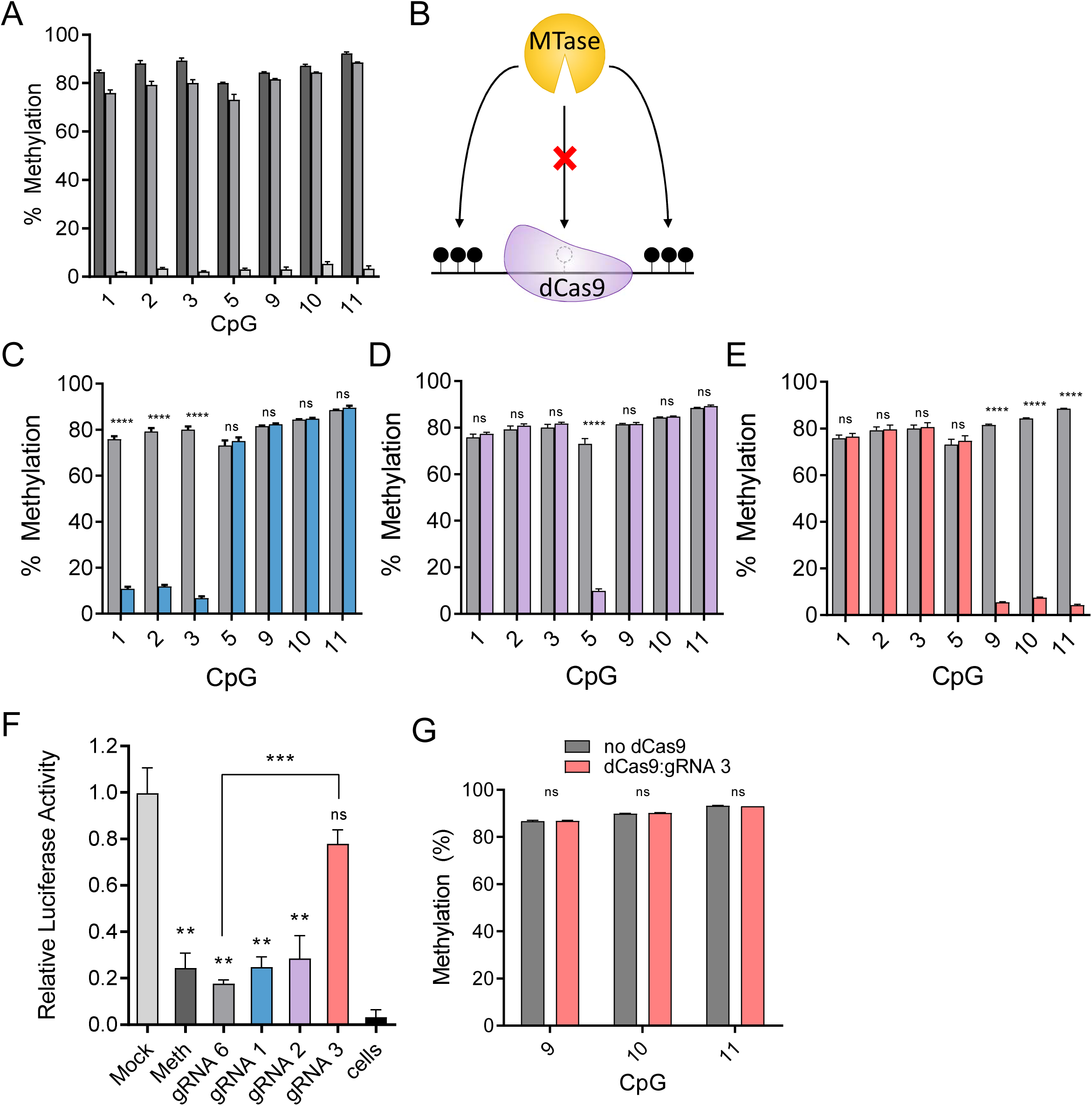
dCas9 blocks DNA methyltransferase *in vitro*. (A) Pyrosequencing data (mean+/-SEM, n=4) for the methylation state of indicated CpGs in the *Il33*-pCpGl plasmid following standard methylation for 4 hours by M.SssI (dark grey), methylation in the presence of dCas9 and gRNA 6 (distant binding) (grey), or a mock-methylated control reaction that lacked S-adenosyl methionine substrate (light grey). (B) Diagram illustrating the principle of site-specific methylation utilizing pre-incubation of DNA with dCas9 and selective CpG-targeting guide restricting M.SssI from binding and methylating the targeted region, while permitting methylation of remaining unobstructed CpGs. (C-E) Pyrosequencing data (mean+/- SEM) for the methylation state of CpGs in the IL-33-pCpGl plasmid following pre-incubation with dCas9 and gRNA1 (C), gRNA2 (D), or gRNA3 (E) and methylation by M.SssI (colored bars). Grey bars are identical in (A, C-E) and indicate methylation levels for the same treatment utilizing gRNA6. (F) Luciferase reporter activity of the plasmids in (A, C-E), expressed as mean+/-SEM relative light units normalized for protein content per sample, and then normalized to average value for mock methylated condition. (G) Percent of methylation (mean+/-SEM) assayed by pyrosequencing when Il33-pCpGl is incubated with dCas9 and gRNA 3 or only gRNA 3 (no dCas9 control) after standard methylation, instead of before (n=3). * indicates statistically significant difference of p<0.05, ** of p<0.01, *** of p<0.001, **** of p<0.0001, and ns = not significant (Student’s t-test, with Holm-Sidak correction if number of tests is greater than 3).

The DNA was then incubated with recombinant dCas9 protein and each of the three *in vitro* transcribed gRNAs – targeting CpGs in the proximal promoter region of *Il33-002* – in order to facilitate binding of the dCas9:gRNA complex to the DNA prior to the addition of M.SssI methyltransferase (Fig. 2B). Following M.SssI treatment, the methylation state of each target CpG was assayed by bisulfite conversion and pyrosequencing and compared to treatment with control gRNA6. Pre-incubation of *Il33*-pCpGl with dCas9 and all CpG-targeting gRNAs resulted in a drastic, specific interference with DNA methylation at targeted sites (Fig. 2C-E). For example, in the case of gRNA3, the targeted CpGs (CpGs 9, 10, and 11) were methylated only to a mean±SEM of 5.75±0.45%, whereas the control gRNA6 barely affected methylation and the sites were methylated at 84.79±0.88% (p<0.00001). Sites that were not directly within or adjacent to the binding site of dCas9:gRNA3 (CpGs 1, 2, 3, and 5) remained unaffected by the treatment (Fig. 2E) (p=0.752, 0.878, 0.800, 0.618, respectively). The same levels of inhibition and specificity were achieved by two other CpG-targeting gRNAs (Fig. 2C and 2D). Notably, with gRNA2, we successfully prevented methylation of a single CpG while leaving all remaining assayed CpGs completely unaffected (Fig. 2D). We also reversed the order of the reaction, incubating the target DNA first with M.SssI and then with dCas9 and gRNA3 in order to ascertain that dCas9 is not able to catalytically remove methyl groups *post hoc* but rather inhibits methylation by competitive binding (Fig. 2G).

Now in possession of five *Il33*-pCpGl plasmids bearing unique methylation patterns (gRNA1, gRNA2, gRNA3, gRNA6, and mock), we sought to assay the impact of these patterns on transcription in live cells using a transient transfection reporter assay. We transfected each uniquely methylated plasmid into NIH-3T3 cells and performed a luciferase reporter assay (Fig. 2F). As mentioned previously, mock (unmethylated) plasmid drove luciferase activity to a significantly higher degree than both standard methylated and dCas9:gRNA6 treated plasmids. When CpGs 1, 2, 3 were unmethylated (by gRNA1 treatment) or CpG5 was unmethylated (by gRNA2), luciferase activity remained low and was not significantly different from gRNA6 control (p=0.202, p=0.332). However, in the case of unmethylated CpGs 9, 10, and 11 (by gRNA3) surrounding the *Il33-002* TSS, luciferase activity was significantly greater than gRNA6 (p=0.0007) and not significantly different from mock-methylated DNA (p=0.157), demonstrating that the methylation of these three TSS CpGs, but not the others, blocks *Il33-002* promoter activity. gRNA1 and gRNA3 both interfered with methylation of 3 CpGs and thus the overall promoter methylation levels were similar between these two treatments; yet, there was a stark difference in luciferase activity. These data demonstrate the exquisite impact of site-specific methylation rather than just methylation density, and thus this assay appears to capture the sequence specificity of inhibition of promoter function by DNA methylation.

In summary, we demonstrate that dCas9 specifically inhibits DNA methylation of targeted sites *in vitro*, enabling the analysis of the causal role of specific methylated sites *per se*. The only difference between our different transfected plasmids is the positions of the methyl moieties. No additional confounding enzyme is introduced. CpGs 9, 10, and 11 at the *Il33-002* TSS silence transcription; demethylation of these CpGs is sufficient for maximal activation of the promoter-reporter construct. In contrast, demethylation of CpGs 1, 2, 3, or 5 is insufficient for re-activation of the methylated promoter suggesting that methylation of these sites is not involved in silencing of transcription from the *Il33-002* promoter.

### Blocking of methylation by dCas9 is limited to its binding site and is symmetrical

In the preceding *in vitro* assays, we were able to prevent on-target DNA methylation with dCas9 without affecting the remaining target CpGs in the promoter. However, as the *Il33-002* promoter is CpG-poor and clusters of CpGs (e.g. 1, 2, 3 and 9, 10, 11) are separated by several hundreds of base pairs, the precision of this approach needs to be determined. In order to delineate the DNA span that is protected from methylation by bound dCas9, we repeated the same *in vitro* assay using a canonical CpG-rich promoter. The human *CDKN2A* (p16) promoter contains a 310 bp fragment with 38 CpGs, which are frequently aberrantly hypermethylated in all common cancers [46]. We designed a gRNA overlapping a single CpG (CpG 17) within this promoter that was flanked on either side by CpGs 8 base pairs away from the 23-nucleotide gRNA and protospacer adjacent motif (PAM) sequence (Fig. 3A). We then applied bisulfite-cloning to map the methylation patterns of individual DNA molecules and assessed whether there was a difference in the methylation pattern of the CpGs in the strand bound by the dCas9:gRNA ribonucleoprotein and its complementary strand (as CpGs are palindromic).

**Figure 3.**
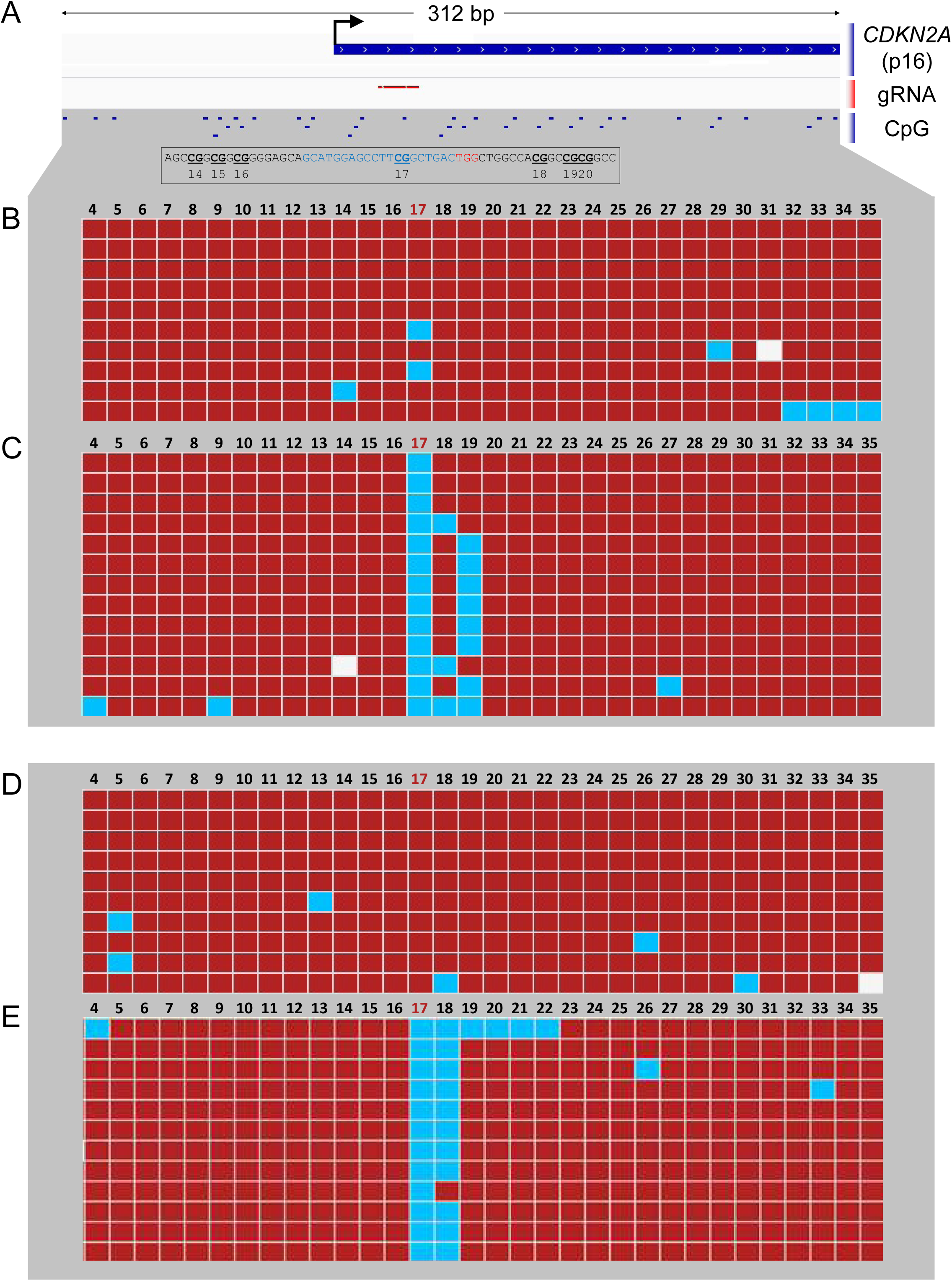
The footprint of dCas9. (A) Genome browser diagram of the *CDKN2A* (p16) promoter region, which was used for the methylation assay, showing transcription start site (TSS, marked by black arrow), gRNA position overlapping CpG 17, and surrounding CpGs. Below, DNA sequence is shown in black, gRNA sequence in blue, and PAM site in red, with CpGs bolded, underlined, and numbered according to the figures that follow. (B-E) Methylation of individual strands of the *CDKN2A* promoter plasmid following standard methylation (B,D) or methylation preceded by incubation with dCas9 and p16 gRNA (C,E). Red squares indicate methylated CpGs and blue squares indicate unmethylated CpGs; white squares indicate no data. Figures (B) and (C) represent the forward strand whereas (D) and (E) represent the reverse strand. Figures generated by BISMA software (http://services.ibc.uni-stuttgart.de/BDPC/BISMA/). Regions below 80% methylation were filtered out as strands that were not effectively methylated by M.SssI.

In the presence of a scrambled control gRNA, M.SssI almost completely methylated all CGs on both strands (Fig. 3B and 3D) with some sporadic unmethylated CpGs that are likely consequences of poor bisulfite conversion or Sanger sequencing errors; M.SssI is highly processive and it is unlikely that the sporadic demethylation resulted from inhibition of M.SssI [47]. In contrast, p16-targeting (CpG 17) gRNA completely inhibited methylation of the targeted CpG on the gRNA bound strand while scrambled control gRNA did not block DNA methylation of CpG 17 (0% vs. 80% methylation, p<0.0001, Fisher’s exact test) (Fig 3B-C). The CpG immediately downstream of the gRNA-PAM sequence was slightly but not significantly unmethylated (77% vs. 100% methylation, p=0.2292, Fisher’s exact test). Interestingly, the following CpG 19 was significantly unmethylated (38% vs. 100% methylation, p=0.0027, Fisher’s exact test), while the CpG only two additional base pairs downstream (CpG 20) was 100% methylated and unaffected. The distance between the unaffected CpG 20 and the 3’ end of the PAM is 14 bp and the upstream unaffected CpG 16 is 8bp from the 5’ end of the gRNA (Fig. 3A). We thus define the range of dCas9 inhibition of M.SssI DNA methylation to be less than 8 base pairs from the 5’ end and smaller than 14 base pairs from the 3’ end of the PAM adding to a total protection range of 45 bp. Nevertheless, peak inhibition is exactly at the binding site and any inhibition within the 45 bp is only partial.

It is interesting to also note that while the target CpG 17 is always protected from methylation in all of the molecules, CpG 18 and/or CpG 19 are protected only in certain DNA molecules. These data suggest that CpGs 3’ of the gRNA sequence are variably protected, possibly reflecting the dynamic orientation of the flexible gRNA scaffold [48]. It may thus be possible to refine this method to reduce or, conversely, target protection of neighboring CpGs. The results are in accordance with the crystal structure of the dCas9:gRNA:DNA triplex, which reveals minimal 5’ protrusion of dCas9:gRNA beyond the 5’ end of the gRNA target and more pronounced extension of both dCas9 protein and, to a larger degree, gRNA scaffold beyond the 3’ end of the gRNA target sequence [48].

We also determined whether protection from methylation by dCas9 was symmetric on both DNA strands and whether dCas9 preferably obstructed methylation of the targeted CpG only on the strand that was complementary to the gRNA. Given that bound dCas9 envelopes nearly the entire DNA double helix [48], we predicted that both CpG sites would be equally protected. Bisulfite-cloning of the opposite strand again revealed complete protection from M.SssI methylation of CpG 17 (0% vs. 100%, p<0.0001) and the next CpG (8% vs. 90%, p=0.0003) (Fig. 2D-E). Interesting, the 3’ footprint is smaller by at least 2bp (and at most 6bp) than in the strand interacting with the gRNA, as CpG 19 is not affected on the antisense strand. Thus, dCas9:gRNA complex completely protected both the target and complementary CpG on the antisense strand.

We determined whether we could focus the range of protection using the smaller dCas9 protein from *Staphylococcus aureus* despite the fact that it requires a longer gRNA (21bp instead of 20bp) and a longer PAM sequence (NNGRR instead of NGG). We designed 4 *S.aureus* gRNAs (SAgRNAs1-4) that also overlapped with potential gRNAs for the hitherto utilized *Streptococcus pyogenes* dCas9 (SPgRNAs1-4) (Supp. Fig. S2A). The first three gRNAs assayed the 5’ protrusion and were shifted by one base pair each in order to refine the 5’ distance both for dCas9 variants; three 5’ CpGs were 4, 7, and 11 bp away from the 5’ end of SAgRNA1; 3, 6, and 10 bp away for SAgRNA2; and 2, 5, and 9 bp away for SAgRNA3. Each CpG was 1 bp further away for the corresponding SPgRNAs as these were 1 bp shorter at the 5’ end (20 bp vs. 21 bp). We determined that *S. aureus* dCas9 is equally capable of complete interference with M.SssI at sites within the bound region (CpGs 20-22), with a gradual 5’ fall-off in protection; 90%-100% protection of CpG 2-4bp away, 80% protection of CpG 5 bp away, 50-60% at CpG 6 or 7 bp away and 0-10% at 9-11bp away from the target (Supp. Fig. S2B and S2C). 5’ interference of SP-dCas9 was consistently less than SA-dCas9 at all distances in a manner that was not sufficiently explained by the additional single 5’ bp of the *S. aureus* gRNAs (Supp. Fig. 2C). The 3’ distance for SP-dCas9 could not be refined further because of a lack of efficacy of SPgRNA 4 (Supp. Fig. S2B and S2D); only 4 strands appeared to have been protected from dCas9 (of 17 sequenced) and the interference was interestingly limited to CpGs in the PAM site and not within the gRNA binding site, likely indicative of a poor-quality gRNA. However, SAgRNA4 was efficient and we could calculate that SAdCas9 interfered with a minimum of 11bp and a maximum of 13bp from the 3’ end, including its 5bp PAM sequence. Therefore, we demonstrate that despite its smaller protein size, SA-dCas9 has a 3’ footprint comparable to but possibly smaller than SP-dCas9 (likely due to similar gRNA scaffolds) and a definitively larger 5’ footprint, drawing the conclusion that the original SP-dCas9 allows more precise interference with DNMTs, however it is also useful to note that the equivalent efficacy of SA-dCas9 presents a secondary option for combinational approaches and for a more diverse selection of target sequences by addition of a second PAM option.

### The dCas9 system directs robust site-specific demethylation in living cells

dCas9 is obviously not an active demethylase; nevertheless, we hypothesized that we could use it to demethylate specific CpGs in living dividing cells. As nascent post-replicative DNA is unmodified and must be methylated by the maintenance methyltransferase DNMT1 in order to preserve parental cell methylation patterns [49], we postulated that dCas9 would interfere with DNMT1 methylation similar to its blockage of M.SssI methylation and thereby cause passive demethylation of targeted sites through successive rounds of cell division and DNA replication. Therefore, we used the gRNAs characterized above to demethylate the endogenous *Il33-002* promoter in NIH-3T3 cells. We established by lentiviral transduction cell lines stably expressing SP-dCas9 and each *Il33* gRNA or a scrambled, non-targeting control gRNA (gRNAscr) and collected DNA for methylation analysis by bisulfite conversion and pyrosequencing one week after complete antibiotic marker selection. We demonstrate that the dCas9:gRNA complex is sufficient to produce robust demethylation of targeted CpGs (Fig. 4 A-C). dCas9 in combination with gRNA1 reduced absolute methylation levels by an average of 27.0% (p<0.0001), 28.3% (p<0.0001), and 34.6% (p<0.0001) at CpGs 1, 2, and 3, respectively; gRNA2 reduced CpG 5 methylation by 52.0% (p<0.0001); gRNA3 reduced CpG 9, 10, and 11 methylation by 30.2%, 31.4%, and 38.4% (p<0.0001 for all). Demethylation with dCas9, unlike dCas9-TET (Fig. 1F) was highly specific to targeted CpGs, as in the case of gRNA2, no other assayed CpGs were demethylated. gRNA3 caused significant demethylation of off-target CpG 3 (p=0.002) but the extent of demethylation was only 0.6%. gRNA1 caused a slightly larger, significant demethylation of the distant CpGs 9, 10, and 11 (5.3%, 4.5%, and 3.9%) but still to a level much reduced in comparison to target CpGs 1, 2, and 3, and less than that of dCas9-TET:gRNA1. These data also clarify that the binding site demethylation in dCas9-TET and in dCas9-deadTET cells (Fig. 1E-G) likely stems from the same mechanism of steric interference with DNMT1 rather than a catalytic TET activity, as the tightly bound dCas9 domain likely makes it impossible for the fused TET domain to access this bound DNA.

**Figure 4.**
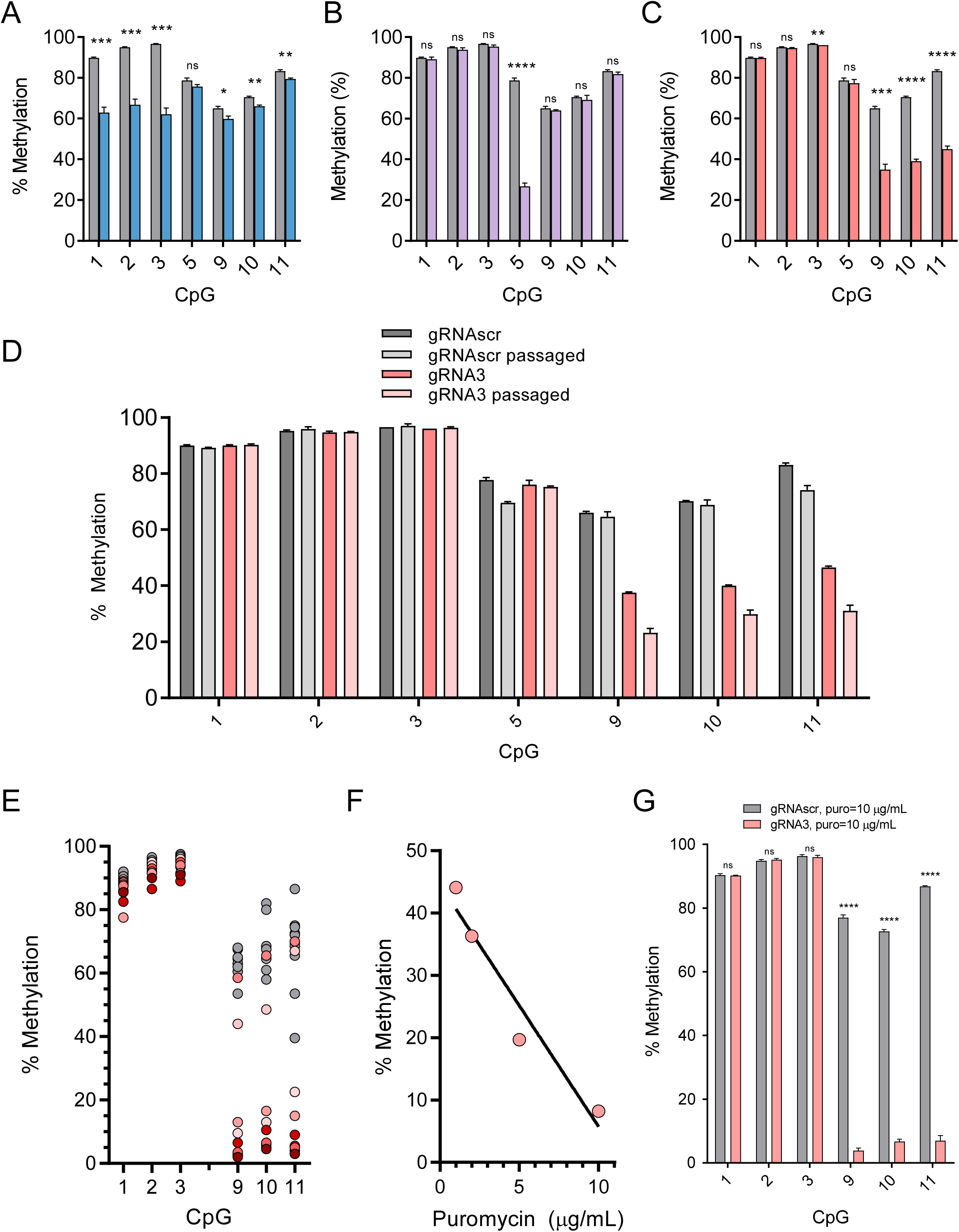
dCas9 causes demethylation in mammalian cells. (A-C) Methylation levels assayed by bisulfite-pyrosequencing at CpGs 1, 2, 3, 5, 9, 10, and 11 of NIH-3T3 cells stably expressing dCas9 and gRNA1 (A, blue), gRNA2 (B, purple), gRNA3 (C, pink) or scrambled gRNA (A-C, grey; identical in all). Data is displayed as mean +/- SEM. (D). Cells from (C) were passaged for an additional 30 days and methylation percentage was assayed as previously (n=3, mean +/- SEM). (E) Cells from (C) were subjected to clonal isolation and expansion. Grey circles represent methylation levels of clones containing dCas9 and scrambled gRNA and various red circles represent methylation levels of randomly selected clones stably expressing dCas9 and gRNA 3 (n=10 per condition). (F) Average DNA methylation at CpGs 9-11, assayed by bisulfite-pyrosequencing, as a function of increasing the selection antibiotic puromycin (lentivirus is expressing puromycin resistance gene) concentration in cell lines (pools) stably expressing dCas9 and gRNA3 (n=1 per puromycin concentration) fitted with a line of best fit. (G). DNA methylation at CpGs 1, 2, 3, 9, 10, and 11 in NIH-3T3 cells stably expressing dCas9 and gRNA3 (pink) or control gRNAscr (grey) and treated with 10 µg/mL puromycin until no antibiotic-associated cell death could be observed and surviving cells were of sufficient quantity for DNA extraction and other procedures (approximately 2 weeks). * indicates statistically significant difference of p<0.05, ** of p<0.01, *** of p<0.001, **** of p<0.0001, and ns = not significant (Student’s t-test, with Holm-Sidak correction if number of tests is greater than 3).

We were also able to demonstrate similar levels of demethylation and specificity by a second gRNA targeting CpGs 9, 10, and 11 which was shifted two base pairs in the 3’ direction (Supp. Fig. S3A) relative to gRNA3, demonstrating that altering the exact CpG positioning relative to the gRNA, whether within the gRNA target sequence, PAM site, or immediately adjacent to either, does not impact demethylation efficiency in cells. All these positions were predicted to be completely protected from DNMT activity by both gRNAs in the *in vitro* footprint assays (Fig. 3).

Though these experiments demonstrated a higher specificity of dCas9 than dCas9-TET across adjacent CpGs in the *Il33-002* promoter, we also sought to determine if the same off-target effects seen with dCas9-TET could be found in equivalent dCas9 treated cells. Unlike in dCas9-TET cells, the distant *Il33-001* transcript was not upregulated by dCas9 combined with any of the three targeting gRNAs (Supp. Fig. S3B); however, there was detectable significant downregulation of *Il33-001* under gRNA1. We also assessed whether dCas9:gRNA3 caused demethylation of the top 5 predicted candidate off-target CpGs for gRNA3 and found that there was no observable change in methylation of any of the top-predicted off-targets (Supp. Fig. S3C, Supp. Table S4).

Next, we wished to evaluate if the dCas9 demethylation approach could be optimized to yield higher demethylation. Passive demethylation by DNMT1 interference would require cell division and if fully efficient, methylation levels would halve with every round of replication. We therefore hypothesized that passaging the cells in culture would increase the extent of demethylation. dCas9:gRNA3 and dCas9:gRNAscr cell lines were passaged for an additional 30 days after the original DNA collection. Indeed, this approach increased the extent of demethylation of only CpGs 9 (14.3%, p=0.0009), 10 (10.2%, p=0.003), and 11 (15.5%, p=0.002) (Fig. 4D). Passaged dCas9:gRNAscr cell lines were demethylated at several CpGs compared to original unpassaged cells but none of these differences were significant after correction for multiple testing.

Another common approach to improve the efficiency of CRISPR/Cas9 editing is cloning [50]. Despite the fact that we could achieve robust demethylation of a target CpG in a population of cells, as a particular strand of DNA only exists in a methylated or unmethylated state, we reasoned that we could isolate clonal populations that are completely demethylated at the target sites (CpG 9,10,11). Therefore, we expanded 10 clonal lines from each of the dCas9:gRNA3 and dCas9:gRNAscr cell lines and subjected these clones to pyrosequencing. The population of gRNAscr clones was not significantly demethylated relative to the original gRNAscr pool at any CpG except a significant 0.6% demethylation at CpG 3, and, with the lone exception of a single CpG in one clone that displayed 39.5% methylation, no CpG in any of the 10 clones was methylated less than 50% (Fig. 4E). Therefore, even though some gRNAscr cells in a population that is not 100% methylated must have fully unmethylated CpGs, the clonal isolation process is unable to generate fully demethylated clones, perhaps due to a given equilibrium between methylation and demethylation established by the nuclear DNA methylation machinery in the cells. dCas9:gRNA3 clones were not significantly demethylated at target CpGs 9, 10, and 11, compared to both original and passaged lines. However, 6 of 10 clones isolated from the dCas9:gRNA3 pool displayed methylation levels below 11% at CpGs 9, 10, and 11 and two of these clones were methylated at or below 5% at all targeted CpGs. We conclude that we were able to produce cell lines with almost completely demethylated target CpGs with this approach (the small level of methylation detected in these clones is around the standard error for unmethylated controls in our pyrosequencing assay).

The clonal analysis suggests a clonal variation in the extent of demethylation by dCas9:gRNA. A plausible cause could be variation in the level of expression of either dCas9 or the gRNA. dCas9 expression did not correlate with methylation levels (Supp. Fig. S3D) whereas gRNA3 expression levels correlated negatively with methylation (r= −0.7307, p<0.05) (Supp. Fig. S3E). Similar to several others studies that demonstrated that expression of gRNA is the rate limiting factor in Cas9 cleavage efficiency [51–54], our data suggest that gRNA is the limiting factor in targeted demethylation efficiency.

Clonal isolation is tedious, involves long passaging times, and prone to producing bottleneck effects; we also found that unhealthy morphologies were common to these clonal populations (Supp. Fig. S4). In order to increase gRNA transgene expression in the clonal population, we increased the quantity of puromycin, which we hypothesized would select for cells with higher copy numbers of virally-inserted transgenes and increased output of gRNA expression. We noted a stepwise increase in demethylation as puromycin concentrations were increased from the standard 1 µL/mL concentration to 2, 5, or 10 µg/mL (Fig. 4F) with a significant correlation (p<0.05) and a large difference of 36% in extent of demethylation of the target sequences between minimal and maximal concentrations. Settling on 10 µg/mL, we produced high-puromycin selected populations of gRNAs1-3 and gRNAscr and verified the extent of demethylation. We found that dCas9:gRNA3-treated cells were highly demethylated at CpGs 9-11 with 3-10% residual methylation, compared to 71-87% in dCas9:gRNAscr cells with 10 µg/mL puromycin (p<0.000001 for all), while off-target CpGs 1-3 were still highly methylated and unaffected by the treatment (p=0.742, 0.621, and 0.670, respectively) (Fig. 4G). In summary, we successfully developed a protocol to produce near-complete, specific targeted DNA demethylation in cell lines and selected this optimized approach for future experiments.

### The effect of site-specific demethylation on *Il33* gene expression

The next step was to assess the utility of our demethylation strategy in exploring the causal links between DNA demethylation at a specific region and transcriptional changes. We predicted that demethylation in this context would not be sufficient to activate transcription because dCas9 remains bound to the TSS and obstructs binding of transcriptional machinery, which is in itself an established technique to inhibit gene expression [40]. Accordingly, despite robust demethylation, high-puromycin dCas9:gRNA3 cell lines expressed significantly less *Il33-002* transcript than even scrambled cells (Supp. Fig. 3F) whereas the *Il33-001* isoform was unaffected in the same cells (Supp. Fig. 3B). In fact, in contrast to the typical negative correlation between expression and DNA methylation, *Il33-002* expression was positively correlated with CpG 9-11 methylation level across dCas9:gRNA3 clones (r=0.74, p=0.02) (Supp. Fig. 3G). This unique relationship likely originates from the fact that increased dCas9 on-target binding not only obstructs DNMT1 activity but also concurrently blocks access to RNApolII complex, inhibiting transcription.

To study the transcriptional consequences of promoter demethylation, dCas9 would need to be removed following demethylation in order to expose the newly unmethylated DNA to the nuclear environment. We tested transient gRNA expression with the aim that following several rounds of cell division, having caused demethylation of target DNA, gRNAs will be diluted and will not block binding of RNApolII. However, transient transfection of guide RNA molecules in a stably expressed dCas9 background resulted in only 15% on-target demethylation (Supp. Fig. S5) and we determined to forego optimization of this strategy in favor of one compatible with the optimized high-puromycin protocol we had established. We implemented the Cre-lox system (Fig. 5A) that would allow complete dCas9 removal by Cre-recombination only after demethylation is maximized.

**Figure 5.**
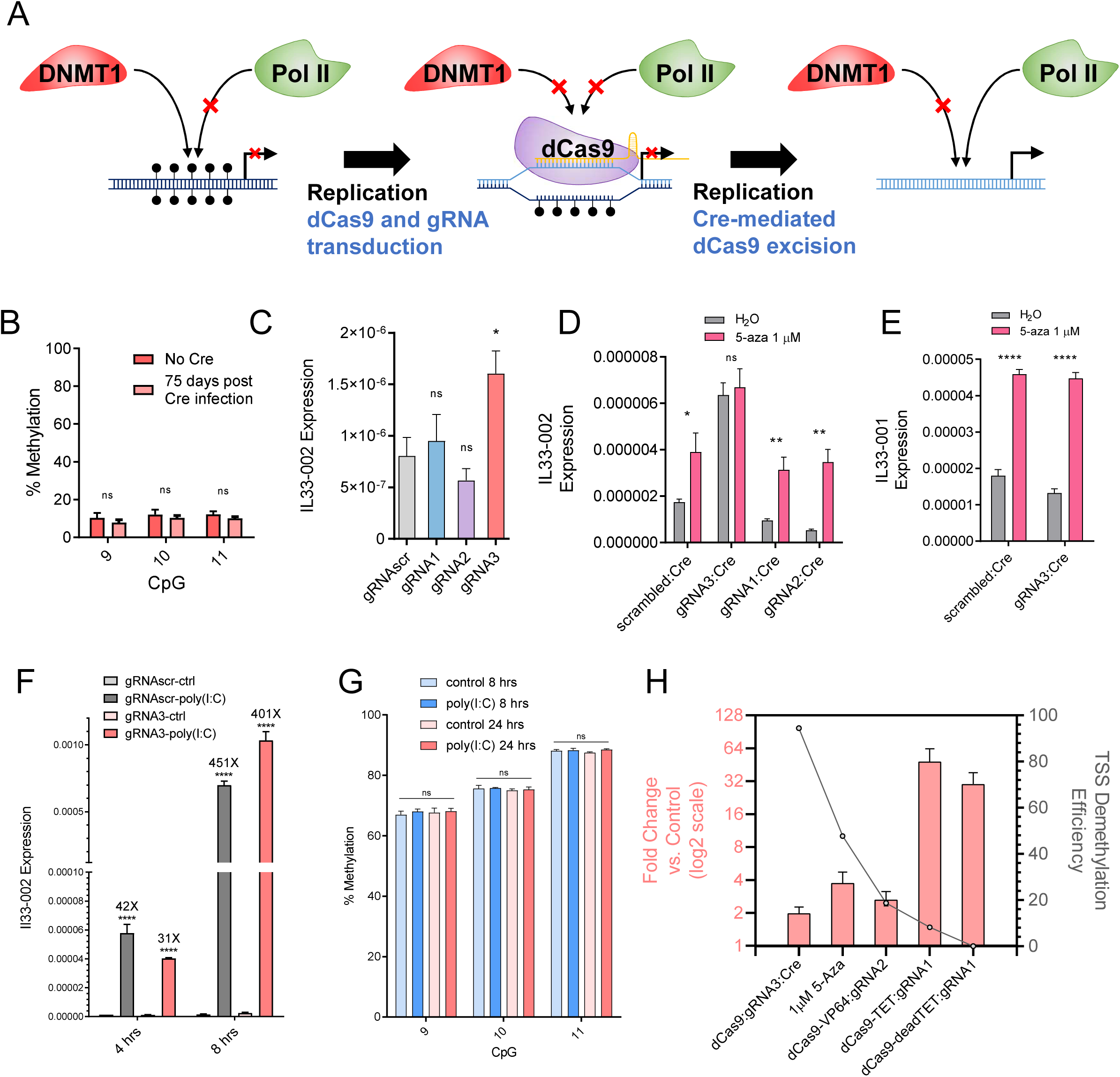
The effect of targeted promoter DNA demethylation on *Il33* expression. (A) Diagram illustrating the principle of site-specific demethylation with dCas9 removal in order to facilitate transcription factor binding to the newly demethylated region. First, DNA is endogenously methylated by DNMT1 with every round of replication and RNA-polII is not recruited to the promoter. After the introduction of dCas9 and a promoter-targeting gRNA, DNMT1 is physically occluded from the locus and nascent strands of DNA are unmethylated, facilitating passive demethylation of the bound region. However, RNA-polII is also physically occluded by dCas9. If dCas9 is successfully removed, the unmethylated DNA no longer serves as a substrate for DNMT1 and continues to remain unmethylated and RNA-polII may now be recruited. (B) Methylation of CpGs 9, 10, and 11 which had been previously demethylated by high-puromycin gRNA3:dCas9 in NIH-3T3 cells, after 75 days of passaging following the lentiviral transduction of Cre recombinase (pink) or empty-vector control (red). (C) *Il33-002* expression in NIH-3T3 cell lines stably expressing gRNAscr (grey) or gRNA1 (blue), gRNA2 (purple), or gRNA3 (pink) under high-puromycin conditions in combination with dCas9, followed by dCas9 removal by Cre recombinase as assayed by RT-qPCR and normalized to *Actb* expression (n=3). (D-E) *Il33-002* expression (D) or *Il33-001* expression (E) in NIH-3T3 cells from (C) following treatment wither water control or 1 µM 5-aza-2’-deoxycytidine, measured by RT-qPCR and normalized to *Actb* expression (n=4-5). (F) Il33-002 expression measured by RT-qPCR and normalized to *Actb* expression, in dCas9:gRNAscr (grey) or dCas9:gRNA3 (pink) NIH-3T3 cells following Cre recombinase treatment and then treated with poly(I:C) (1 µg/mL) or water control for 4 or 8 hours. (G) DNA methylation assayed by bisulfite-pyrosequencing in NIH-3T3 cells expressing dCas9, gRNAscr, and Cre treated with 1 µg/mL poly(I:C) or water control for 8 hrs and 24 hrs (n=3). (H) Maximal Il33-002 induction (left y-axis, pink bars; data in log2 scale but axis numbering is not transformed) and maximal promoter demethylation (right y-axis, calculated as percent unmethylated divided by control methylation) under different treatments (x-axis: dCas9, 5-aza-2’-deoxycytidine, dCas9-VP64, dCas9-TET, and dCas9-deadTET). Where relevant, data for maximally inducing/demethylating gRNA is shown. * indicates statistically significant difference of p<0.05, ** of p<0.01, *** of p<0.001, **** of p<0.0001, and ns = not significant (Student’s t-test, with Holm-Sidak correction if number of tests is greater than 3).

We established new high-puromycin selected NIH-3T3 cell lines expressing each lentiviral *Il33* gRNA and a lentiviral loxP-flanked dCas9 variant and validated successful demethylation (Supp. Fig S6A-C). One of the two base substitutions to render this dCas9 variant nuclease-dead (D10A, H840A) is different than the dCas9 used in previous experiments (D10A, N863A). We then used lentivirus-mediated gene transfer to introduce either Cre recombinase or an empty control vector and verified successful dCas9 removal by Cre at the DNA level by PCR, using primers that produced a 500bp fragment upon recombination (Supp. Fig. S6D-E), and at the protein level by chromatin immunoprecipitation followed by quantitative PCR (ChIP-qPCR). ChIP-qPCR demonstrated elevated dCas9 binding to the *Il33-002* promoter region only in cells stably expressing dCas9 and gRNA3 but not in dCas9:gRNAscr cells regardless of Cre treatment (Supp. Fig. S6F); in dCas9:gRNA3 cells, Cre recombination eliminated dCas9 binding to the *Il33-002* promoter. Interestingly, low levels of methylation persisted for at least 75 days after removal of dCas9 by Cre recombinase (Fig. 5B), indicating a lack of *de novo* methylation of this locus in these cells and the ability of this approach to modify DNA methylation in a stable manner despite elimination of dCas9.

Having generated NIH-3T3 cells bearing highly effective targeted demethylation without bound dCas9 to hinder RNApolII binding to the TSS, we were then able to interrogate whether demethylation of the proximal promoter causes changes in expression of the gene. Expression levels of *Il33-002* transcript were measured by RT-qPCR. We detected a small but significant (p=0.0312) increase in expression in NIH-3T3 cells treated with dCas9:gRNA3 and Cre recombinase as compared to dCas9:gRNAscr, but not in dCas9:gRNA1 or dCas9:gRNA2 cells (Figure 5C). This is consistent with our *in vitro/*transient transfection luciferase assays findings (Fig 2F); both approaches suggest that methylation of TSS CpGs 9, 10, and 11 silence the basal *Il33-002* promoter.

It is possible that the small magnitude of induction of expression by demethylation of the TSS region can be explained by the presence of other methylated regulatory regions or other required *trans*-acting factors that need to be demethylated to facilitate larger changes in expression. We used 5-aza-2’-deoxycytidine, a global demethylation agent, to assess whether demethylation of other sites would further induce the expression of TSS-demethylated *Il33-002*. Our results show that gRNAscr-, gRNA1-, and gRNA2-bearing cells, which were still methylated at the TSS, were still induced by the drug, while gRNA3 treated cells that were demethylated at the TSS were no longer responsive (Fig 5D.), suggesting that no further demethylation is required beyond demethylation of TSS sites 9, 10, and 11 for the activity of the basal promoter. To further corroborate that the lack of further induction by 5-aza-2’-deoxycytidine in cells with demethylated CpG sites 9, 10, and 11 was not a consequence some other resistance to demethylation of dCas9:gRNA3 cells, we demonstrate that, in these dCas9:gRNA3 cells, the induction of the *Il33-001* isoform, driven by an untargeted upstream promoter, continued to be responsive to 5-aza-2’-deoxycytidine (Fig. 5E).

We verified that lack of further induction of gRNA3 demethylated *Il33-002* by a demethylating agent was not a result of an upper threshold of expression or our detection method, because treatment of cells with 1 µg/mL polyinosinic:polycytidylic acid (poly(I:C)) activated expression of *Il33-002* several hundred-fold after 4 and 8 hrs (Fig. 5F). Interestingly, this strong induction in response to poly(I:C) occurred in the complete absence of any detectable demethylation of the three TSS CpGs after 8 hours and even when incubation was extended to 24 hours (Fig. 5G) nor of any other CpGs in the promoter (Supp. Fig. S7A). These data suggest that DNA methylation suppresses basal activity of the *Il33-002* promoter but does not affect its inducibility, which can be independent of DNA methylation in the promoter region.

Histone deacetylase inhibition has been previously reported to act in combination with DNA demethylation to activate gene expression [55]. Activation of gene expression might require both demethylation and histone acetylation. We tested whether we can achieve a robust activation of the demethylated *Il33-002* with the histone deacetylase inhibitor trichostatin A (TSA, 50 nM). However, we only noticed a minor difference in the responses to treatment with TSA: in gRNAscr, gRNA1, and gRNA2 cells, TSA slightly reduced expression, and in gRNA3 cells, expression was not affected by TSA (Supp. Fig. S7B). Thus, TSA inhibition of histone deacetylase activity does not add to the transcription activity of *Il33-002*. Finally, we determined whether demethylation poises *Il33-002* to activation by other inducers. LPS has been previously been shown to induce *Il33* [56]. Treatment of NIH-3T3 cells with lipopolysaccharide (LPS, 100ng/mL) induce the expression levels of *Il33-002* in gRNA3 cells where the TSS is demethylated to a larger extent than in cells were *Il33-002* TSS is methylated (gRNAscr), however the fold change relative to phosphate-buffered saline control was similar (Supp. Fig S7C). This suggests that LPS can activate both the unmethylated and methylated *Il33-002*, but the total output increases once the promoter is demethylated. Alternatively, since *Il33* is not 100% methylated in control cells and in some cells the promoter is unmethylated (∼20%), LPS might have induced the unmethylated copies in the control cells explaining the lower total output in the control cells. However, the ratio of unmethylated Il33 promoter (20%) in the untreated cells relative to the demethylated cells (90%) (0.22) is lower than the ratio of expression in control and demethylated cells following LPS induction (0.5). These data are consistent with the hypothesis that methylation silences basal promoter activity but does not affect inducibility.

In summary, we show that near-complete demethylation of *Il33*-*002* using an enzyme-free approach results in only a mild two-fold induction of basal gene expression, whereas other approaches that cause smaller degrees of demethylation can produce larger changes in gene expression, such as dCas9-TET:gRNA1, which produces only a 10% demethylation but a 50-fold gene induction (Fig. 5H). As the larger magnitude of demethylation observed in the dCas9 approach does not produce such substantial transcriptional changes, it is clear that promiscuous mammalian enzymatic domains do not exclusively demethylate, have other methylation independent activities, and cannot be suitably applied to investigate the causational relationship between DNA methylation at specific sites and gene expression.

### dCas9-based demethylation analysis of the role of TSS methylation in *SERPINB5*, *Tnf* and *FMR1* genes

Our previous results show that methylation of *Il33-002* TSS silences basal promoter activity but that demethylation does not result in robust activation of the gene. Induction of this gene could occur independently of methylation of the promoter. We therefore examined whether TSS (de)methylation might play similar or different roles in other genes.

We next examined the *SERPINB5* gene, which encodes the tumor suppressor Maspin and is methylated and transcriptionally silenced in human MDA-MB-231 breast cancer cells. Reactivation of this gene has been reported to increase cell adhesion and therefore decrease growth, invasion, and angiogenesis [57–61]. Several studies have reported that DNA methylation of the *SERPINB5* promoter negatively correlated with gene expression in human cancer and that 5-aza-2’-deoxycytidine treatment is sufficient to restore *SERPINB5* expression [62–66].

We designed a single gRNA targeting 6 CpGs (3 within the gRNA binding site and 3 within 11bp of the 3’ end of the gRNA, as predicted to be completely affected by our *in vitro* footprint assays in Fig. 3) in the core *SERPINB5* promoter and specifically in the transcription-regulatory GC-box (Fig. 6A). In this case, increasing puromycin had a mild effect in increasing the frequency of unmethylated promoters and even the highest puromycin concentrations (40 µg/mL) resulted in demethylation of only 20% (Supp. Fig S8). Therefore, we turned to the previously described clonal isolation strategy. We picked approximately 20 clones from each of the two treatments (gRNAscr and gRNAmaspin) and evaluated methylation by pyrosequencing, which revealed a significant demethylation in gRNAmaspin MDA-MB-231 clones on average comparted to gRNAscr clones (Fig. 6B). We found that numerous clones were completely demethylated by the treatment (Fig. 6C) and we selected 5 gRNAmaspin clones with methylation levels below 5% at all six CpGs as well as 5 representative gRNAscr clones. Surprisingly, despite the large change in methylation, *SERPINB5* expression after Cre-mediated dCas9 removal remained unchanged between the two sets of clones, though there was a small insignificant (p=0.105) increase in the variance expression in the different demethylated clones (Fig. 6D). The difference in *SERPINB5* expression was increased when these cells were further subcloned (Fig. 6E), but not to statistically significant degree (p=0.0767), suggesting that demethylation of the *SERPINB5* promoter is insufficient to activate the gene. Since 5-aza-2’-deoxycytidine was shown to induce the gene we tested whether induction of the gene requires additional demethylation beyond the gene TSS: we tested whether 5-aza-2’-deoxycytidine would induce the methylated and unmethylated *SERPINB5* promoter to the same extent. In contrast to *Il33-002*, which was not further induced by 5-aza-2’-deoxycytidine after TSS demethylation, expression of *SERPINB5* with a demethylated TSS region was significantly increased by 5-aza-2’-deoxycytidine treatment as compared to gRNAscr cells treated with 5-aza-2’-deoxycytidine (Fig. 6F) (p=0.0184) (4.85X in gRNAmaspin vs. 2.59X in gRNAscr). This is consistent with the conclusion that demethylation of the promoter is insufficient for its expression and demethylation of other regions, such as the depicted enhancer regions (Fig. 6A), is required for induction of *SERPINB5*; however, basal promoter demethylation contributes to the overall expression level following demethylation of other regions.

**Figure 6.**
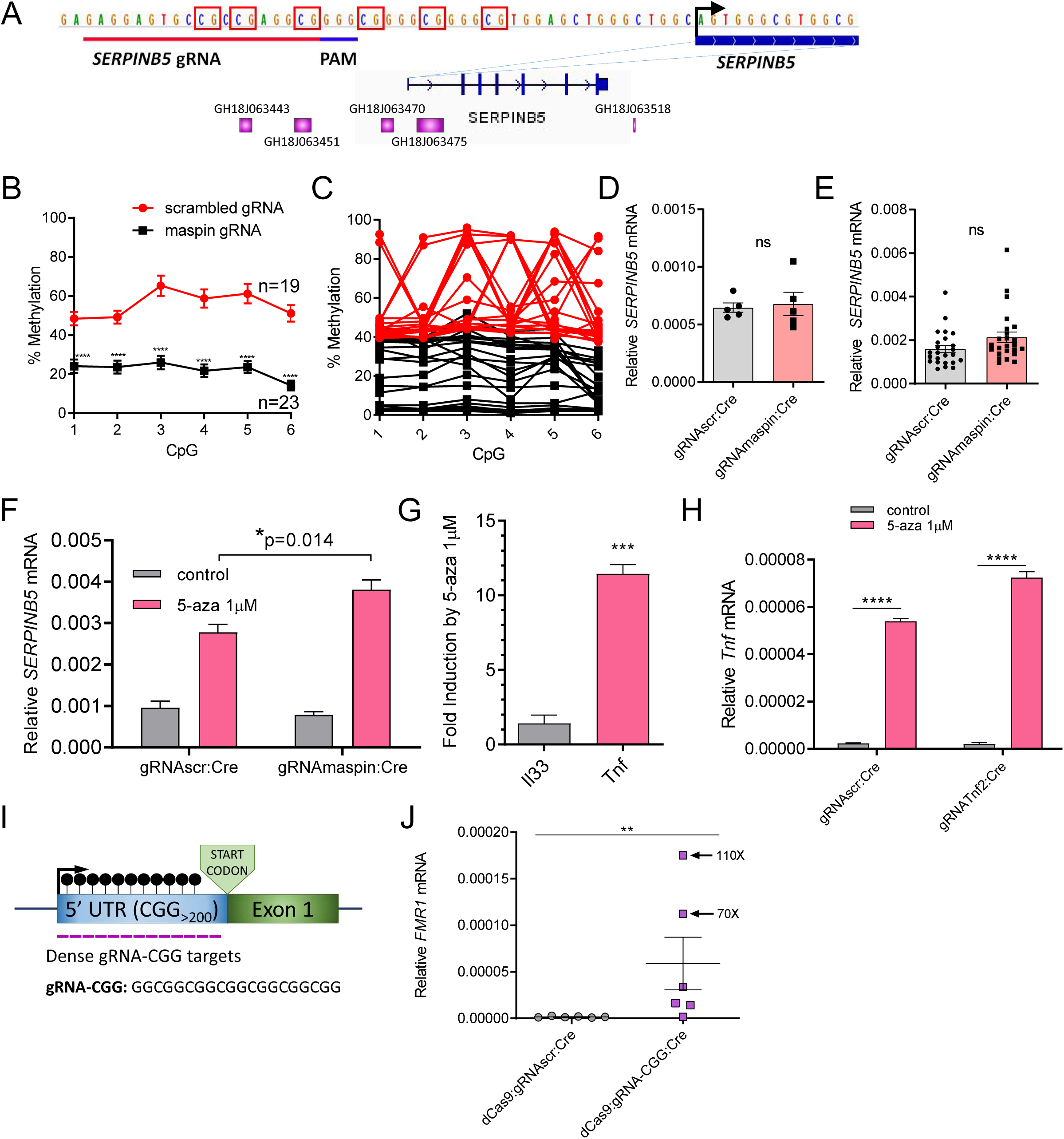
The effect of dCas9-based demethylation of TSS on expression of *SerpinB5*, *Tnf* and *FMR1* genes. (A) (Top) Schematic of the human *SERPINB5* promoter region, including the start site of transcription (marked by black arrow) and the binding site and PAM of the *SERPINB5* gRNA. CG sequences are boxed in red. (Bottom) *SERPINB5* gene with purple boxes indicating enhancer positions relative to gene body. Enhancer IDs correspond to the GeneHancer database. (B) DNA methylation level of each CpG averaged over 19 gRNAscr (red) and 23 gRNAmaspin (black) MDA-MB-231 clones as assessed by pyrosequencing (mean ± SEM). (C) Same data as (B) except now shown as the calculated methylation fraction for each of the 19 gRNAscr (red) and 23 gRNAmaspin (black) clones, rather than the average of all clones. (D) *SERPINB5* expression levels measured by RT-qPCR and normalized to *GAPDH* expression levels for 5 gRNAscr and 5 lowly-methylated gRNAmaspin clones (mean ± SEM, n=5). (E) *SERPINB5* expression levels measured by RT-qPCR and normalized to *GAPDH* expression levels for 48 (24 for each treatment) MDA-MB-231 clones subcloned from the clones in (D). (F) *SERPINB5* expression levels measured by RT-qPCR and normalized to *GAPDH* expression levels for clones from (D) following treatment with 1 µM 5-aza-2’-deoxycytidine or water control (n=5). (G) Expression fold change of murine *Il33-002* (grey) and *Tnf* (pink), normalized to *Actb* and water control, following treatment of control NIH-3T3 cells with 1 µM 5-aza-2’-deoxycytidine (n=3). (H) *Tnf* expression in NIH-3T3 cell lines stably in control (water); grey bars) or 1 µM 5-aza-2’-deoxycytidine (pink bars) expressing either gRNAscr or gRNA*Tnf2*:Cre under high-puromycin conditions in combination with dCas9, followed by dCas9 removal by Cre recombinase, as assayed by RT-qPCR and normalized to *Actb* expression (n=3). (I) Schematic of the human *FMR1* repeat region showing the 5’ untranslated region (UTR) that is prone to CGG repeat expansion and methylation in Fragile X syndrome. Sequence of the gRNA targeting this region is shown (gRNA-CGG) and the extent of the available binding sites for this gRNA is represented by purple lines which indicate binding sites, the 13 presented here represent less than 15% of the available binding site in the Fragile X syndrome patient primary fibroblasts used in this study, which have approximately 700 CGG repeats. (J) *FMR1* expression quantified by RT-qPCR and normalized to *GAPDH* expression levels in Fragile X syndrome patient primary fibroblasts that had stably expressed dCas9 (later removed with Cre) and either gRNAscr (grey) or gRNA-CGG (purple) under high-puromycin selection (n=6). * indicates statistically significant difference of p<0.05, ** of p<0.01, *** of p<0.001, **** of p<0.0001, and ns = not significant (Student’s t-test, with Holm-Sidak correction if number of tests is greater than 3). Exceptionally, for (J) Mann-Whitney U test was used due to unequal variance.

We then questioned whether larger changes in expression could follow demethylation of proximal promoters in other genes. To identify genes that may potentially display such changes, we selected 17 candidate genes in NIH-3T3 cells with large expression fold changes in response to 5-aza-2’-deoxycytidine in a publicly available microarray dataset (GEO GSE8374) and analyzed their expression changes by RT-qPCR following 1µM 5-aza-2’-deoxycytidine treatment (Supp. Table S5). We selected the *Tnf* gene which was heavily methylated at the proximal promoter region and the expression of which was increased by more than ten-fold by 5-aza-2’-deoxycytidine treatment. We tested six gRNAs under high-puromycin selection (20 µg/mL) conditions and identified a gRNA that demethylated all 10 CpGs in approximately 200bp upstream of the *Tnf* TSS (Supp. Fig. S9A-C). We chose this gRNA (gRNATnf2) for Cre recombinase removal of dCas9. Surprisingly, complete *Tnf* promoter demethylation did not result in a significant difference in *Tnf* expression compared to gRNAscr (Fig. 6H) nor could we observe any difference in expression in subclones from these cell pools (Supp. Fig. S8D). However, when these cells were treated with 5-aza-2’-deoxycytidine, the demethylated gRNA*Tnf2* cells were induced to a larger extent than the methylated gRNAscr pools (36-fold versus 24-fold) (p=0.0008) (Fig. 6H). Therefore, we conclude that, similar to demethylation of *SERPINB5* TSS, demethylation of *Tnf* basal promoter contributes to expression but is insufficient to induce expression and that expression necessitates demethylation of a different region either in *cis*, such as the two murine proximal *Tnf* enhancers (Fig. S9A) [67], or in *trans* through activation of putative transcription factors.

Our final demethylation target was the *FMR1* gene which, in patients with Fragile X syndrome, undergoes a CGG repeat expansion (>200 repeats) in its 5’ UTR that becomes aberrantly hypermethylated and results in silencing of *FMR1* transcription [68]. The CGG repeat expansion is a unique target for a guide RNA with the sequence GGCGGCGGCGGCGGCGGCGG and PAM motif CGG since it should bind sequentially to the entire large repeat region and – under sufficient expression levels – shield the entire region from methyltransferase activity (Fig. 6I). We obtained primary fibroblasts from a patient with Fragile X syndrome with approximately 700 CGG repeats exhibiting high methylation [69] – and a lentiviral vector bearing the CGG-targeting gRNA sequence (gRNA-CGG) [70]. After application of our optimized dCas9-demethylation protocol using gRNA-CGG or gRNAscr (20 µg/mL puromycin) we measured expression levels of the *FMR1* gene in 6 independent pools and found significant upregulation of *FMR1* gene expression (p=0.0087) up to a maximum of 110-fold in one pool, as well as other pools with 9-, 10-, 21-, and 70-fold inductions (Fig. 6J), all of which are vastly higher than the induction following TSS demethylation observed in *Il33* and suggestive of the fact that in this case DNA methylation of the repeat region has a large effect on gene expression. We were unable to map DNA methylation in the dCas9 demethylated pools due to technical challenges in amplification of such a highly repetitive and CG-dense region, despite attempting to do so with buffers and polymerases optimized for GC-rich templates, as well as bisulfite-free approaches in an effort to avoid the decreased sequence diversity caused by sodium bisulfite (C to T transitions).

In summary, we demonstrate that the dCas9 demethylation method can be effectively applied in a number of different cell types: a murine fibroblast cell line, a human breast cancer cell line, and primary patient fibroblasts and across different genetic contexts. This method could be used to assess the relative contribution of DNA methylation in specific sites to modulation of gene expression and to delineate positions whose demethylation would have the largest effect on expression. Since our method physically targets DNA methylation without confounding enzymatic activities it provides an unconfounded and at times surprising assessment of the role of DNA methylation.

### CRISPR/Cas9-induced demethylation confounds mutational studies with Cas9

The catalytically active CRISPR/Cas9 system has become the gold standard technique for generating gene knockouts in functional studies. A common technical consideration in these approaches is to target 5’ constitutive exons such that frameshift mutations are more likely to take effect early and render the translated protein nonfunctional [71]. This inevitably results in the positioning of the Cas9:gRNA ribonucleoprotein complex near the TSS and proximal promoter of the targeted gene. Based on the results describe here, we hypothesized that the residence time of DNMT-interfering Cas9, in addition to the drastic epigenetic changes that occur during post-mutagenesis repair [72], may in certain cell subpopulations result in DNA demethylation and gene induction that would confound the interpretation of the results.

We had in a previous study used Cas9 and an *HNF4A*-targeting gRNA from the commonly used GECKO gRNA library [71] to generate *HNF4A* gene knockouts in primary human hepatocytes [73]. The gRNA target site is located in the first exon of several *HNF4A* isoforms, the *HNF4A* TSS is only 2 bp from the 3’ end of the PAM, and there are 3 CpGs directly within the site, with two additional CpGs in close proximity (Fig. 7A). We analyzed one mixed *HNF4A* CRISPR:Cas9 targeted cell population and mapped by Sanger sequencing different *HNF4A* alleles, which were primarily bearing a T->C missense mutation as well as in-frame and out-of-frame deletions (Fig. 7A-C), indicating that a considerable fraction of cells in this population were likely to produce a protein that retained some degree of functionality. To our surprise, we found that this highly methylated region was completely demethylated in this cell population, irrespective of the mutation induced by Cas9 (Fig. 7D). This demethylation was not only extremely substantial but also extremely broad, covering not just a 311bp fragment with 15 CpGs highly methylated in gRNAscr cells to over 90% on average, but also continued to a slightly smaller degree into originally less methylated regions immediately upstream (230bp with 5 CpGs) and downstream (269bp with 7 CpGs) (Fig. 7E). We also found that this demethylated gHNF4A population expressed approximately 15-fold more *HNF4A* mRNA than gRNAscr controls (Fig. 7F). Thus, standard CRISPR/Cas9 gene depletion studies might be confounded by the effects of extensive demethylation.

**Figure 7.**
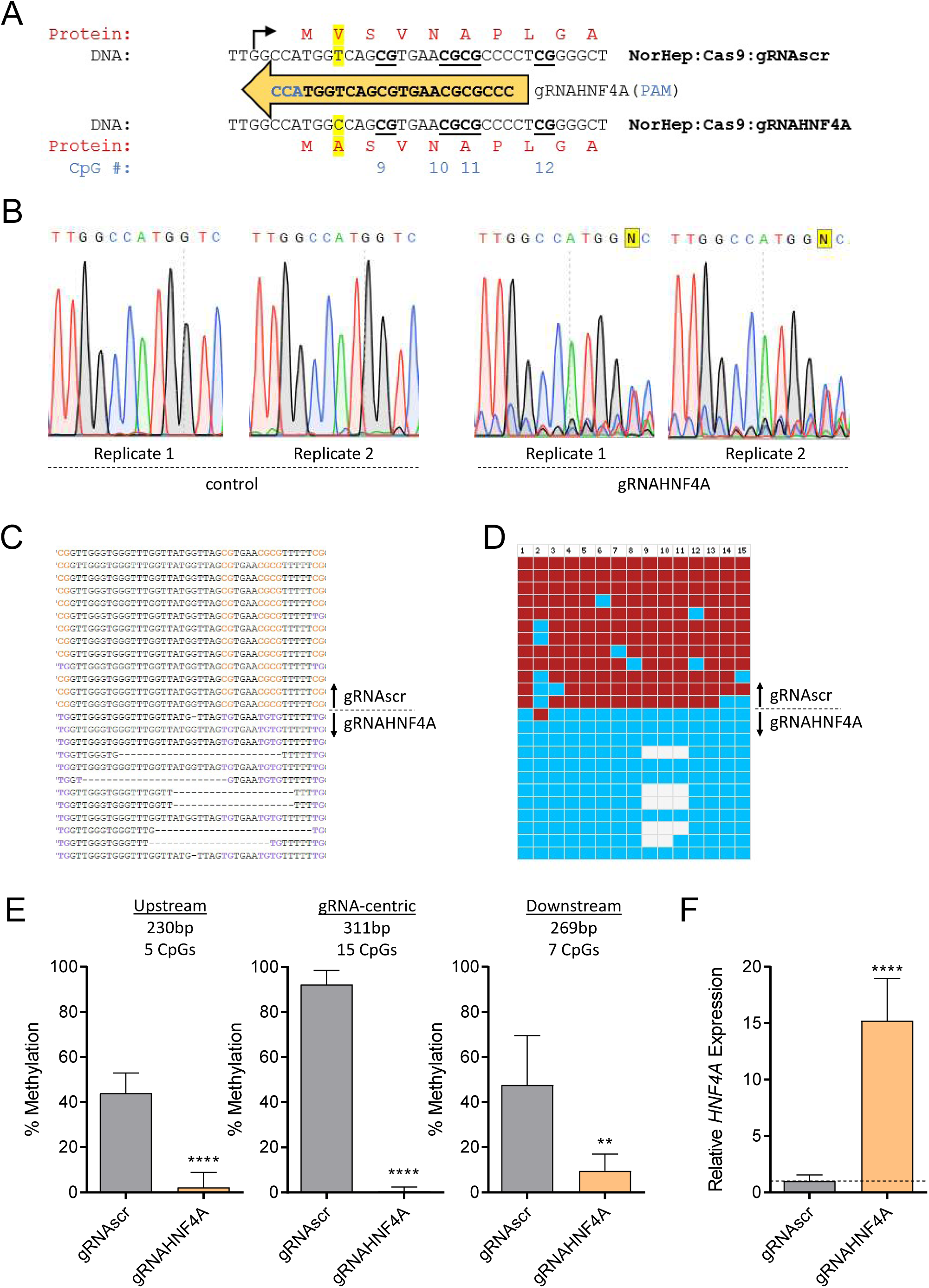
Demethylation is a confound of Cas9 knockout gene deletion. (A) The sequence of the lentiviral gRNAHNF4A and its PAM site in blue. Above is the reference sequence of the *HNF4A* gene near the gRNA target site, as validated by Sanger sequencing in primary human hepatocytes expressing via lentivirus Cas9 and gRNAscr, with the TSS indicated by a black arrow and the reference protein sequence in red. CGs are bolded and underlined. Below is the dominant Sanger sequence profile of a primary human hepatocyte population expressing lentiviral Cas9 and gRNAHNF4A. This mutation and the resulting difference in the amino acid sequence, as well as the reference sequences at this location, are highlighted in yellow. (B) Two technical replicates each of the Sanger sequencing chromatograms from the primary human hepatocytes expressing dCas9 and gRNAscr (left) or dCas9 and gRNAHNF4A (right) at the targeted *HNF4A* locus. (C) Sanger sequencing results of 13 gRNAscr and 12 gRNAHNF4A DNA strands following bisulfite conversion from the cell populations in (B), demonstrating both the methylation levels and the variety of mutations induced by Cas9 in gHNF4A-treated cells. (D) Same as (C) except data expanded is expanded to a larger (>300bp) region, and simplified such that only CpGs are shown, where blue squares indicate unmethylated CpGs, red squares indicate methylated CpGs, and white squares indicate missing information due to Cas9-induced deletions. CpGs are numbered in accordance with (A). (E) Bisulfite-sequencing data from (D) (center) as well as 5 CpGs immediately upstream (left) and 7 CpGs immediately downstream (right), displayed as percent DNA methylation over all sequenced DNA strands in primary human hepatocytes expressing Cas9 and either gRNAscr (grey) or gRNAHNF4A (orange) and as mean ± SD. (F) *HNF4A* expression in primary human hepatocytes expressing Cas9 and either gRNAscr (grey) or gRNAHNF4A (orange) quantified by RT-qPCR and normalized to *GAPDH* expression, followed by normalization to average expression in gRNAscr cells, with a dashed line at 1 (n=6, mean ± SD). * indicates statistically significant difference of p<0.05, ** of p<0.01, *** of p<0.001, **** of p<0.0001, and ns = not significant (Student’s t-test, with Holm-Sidak correction if number of tests is greater than 3).

## Discussion

The extremely consistent developmental profiles of DNA methylation across every human tissue [74] combined with the fact that deviations from these patterns are consistent indicators of disease [1, 75, 76] suggest that DNA methylation has an important role in physiological processes. Importantly, it has been suggested that DNA methylation plays a functional role in the molecular pathology of cancer [1, 3, 64, 76–78] and other common diseases, including mental health disorders [79–81].

Correlation studies since the early 1980s have suggested that DNA methylation in promoters and other transcriptional regulatory regions is negatively correlated with gene expression [82–85]. In the last three decades, several lines of evidence have provided support to the causal role of DNA methylation in the modulation of gene expression. First, *in vitro* methylation of reporter plasmids was shown to silence transcriptional activity when these plasmids were transfected into cell lines [84]. Later studies used different methods to limit *in vitro* methylation to specific regions. Although these studies provide the most direct evidence that there are cellular mechanisms to recognize DNA methylation in particular regions and translate this into silencing of gene activity, the main limitation of these studies is that silencing of ectopically methylated DNA might not reflect on genomic methylated sites and might instead represent a defense mechanism to silence invading viral and retroviral DNA [86] rather than a mechanism for cell-type-specific differential gene expression. Second, DNA methylation inhibitors 5-aza-2’-deoxycytidine and 5-azacytidine provided early evidence for a causal role for DNA methylation in defining cellular identity and cell-type-specific gene expression [87]. However, these inhibitors act on DNA methylation across the genome and do not provide evidence for the causal role of methylation in specific regions or specific genes. Moreover 5-azacytidine was reported to have toxic effects unrelated to DNA methylation [12]. Antisense [13], siRNA[14] and gene knockout [15] depletions of DNA methyltransferases (DNMTs) provided further evidence for the role of DNA methylation in cellular differentiation and development, however DNMT depletion similarly reduces methylation in a general manner, leaving unanswered questions as to the relative role of DNA methylation at specific regions. Furthermore, all DNMTs form complexes with chromatin silencing proteins and might control gene expression by DNA methylation independent mechanisms [77, 88–90].

A study examining the state of methylation of TSS regions that are physically engaged in transcription using ChIP-sequencing with antibody against RNAPolII-PS5, the form of RNApolII that is engaged at transcription turn on, showed that promoters that are actively engaged in transcription onset are devoid of methylation [91]. Although these data show that transcription initiation is inconsistent with DNA methylation, the question of causality remains: is DNA demethylation a cause or effect of transcription onset? Similarly, enhancers are demethylated at transcription factor binding sites; is demethylation a cause or effect of transcription factor binding [92–94]?

To address this longstanding question, CRISPR/Cas9 fusion constructs with TET catalytic domains were generated to target demethylation to specific regions and to determine whether demethylation of particular regions alters transcription activity [19, 20, 95].

Here, we show that while dCas9-TET induces only modest demethylation of the TSS, it induces robust activation of the *Il33-002* gene (Fig. 1), but the results leave us with unanswered questions on whether DNA demethylation of the basal promoter was causal for this activation. First, TET enzymes are not enzymatically demethylases but monooxygenases which oxidize 5-methylcytosine to 5-hydroxymethylcytosine, 5-formylcytosine, and 5-carboxylcytosine, which have demonstrated stability [43, 44], demonstrated differential protein interactors [21–26], and demonstrated structural effects on DNA [27], suggesting that each derivative may be a unique epigenetic mark in its own right. We show here that dCas9-TET causes hydroxymethylation of the *Il33-002* promoter that is maintained in culture (Supp. Fig. S1G). Moreover, TET proteins are also able to oxidize thymine to 5-hydroxymethyluracil, thereby introducing another confounding epigenetic mark that produces a unique spectrum of modifications on chromatin structure and transcription factor activity [96]. A recent candidate for the improvement of such a strategy is the fusion of dCas9 to the *Arabidopsis* ROS1 glycosylase that directly removes 5-methylcytosine by direct base excision repair, foregoing the intermediate oxidized derivatives with epigenetic potential [97]; yet the issue of the overexpression of an enzyme with a capacity for unwanted and non-targeted effects is not solved by this approach.

Moreover, our data suggest that TET activation of *Il33-002* is independent of DNA demethylation since a dCas9-deadTET mutant with inhibited catalytic monoxygenase activity does not trigger demethylation but also activates *Il33-002* to a similar extent as the catalytically active dCas9-TET (Fig. 1H). We also find that the TET2 catalytic domain is capable of inducing unmethylated DNA (Fig. 1J), clearly indicating a demethylation-independent transactivation capacity. It is indeed known that even the restricted catalytic TET domains used in dCas9-TET fusions retain a protein interaction domain that binds O-linked N-acetylglucosamine transferase (OGT) [28, 29] and TET proteins and OGT have been shown to co-localize across the genome [98]. The recruited OGT regulates gene expression by glycosylating and modulating the activity of transcription factors such as HCFC1, SP1, OCT4, MYC, p53, and RNA polymerase II as well as histones to directly increase local H2B mono-ubiquitination and trimethylation of histone 3 on lysine 4, both of which are associated with increased gene expression [28, 98, 99]. This mechanism as well as other potential mechanisms of catalytic-independent transcriptional activation by TET may explain our observation that dCas9-deadTET led to substantial gene induction despite an apparent lack of catalytic activity. The fact that catalytically dead TET protein activates transcription is consistent with previous reports [100] and confounds the interpretation of the causal role of TET induced demethylation in gene activation.

Third, the fact that an enzyme with such a potential for transcriptional modulation is being overexpressed as a dCas9-TET1 fusion introduces capacity for unwanted transcriptional changes, and more recent attempts to use the SunTag system to amplify TET binding at a desired locus [20] only aggravate this issue by overexpression of large numbers of antibody-fused TET1. These undesirable effects would only be negligible in a scenario where a cell expresses a single copy of dCas9-TET that is bound at the intended locus, with highly effective oxidation and base excision repair, an impossible situation given that these lowly-active fusions must be highly expressed to facilitate robust demethylation, and thus inevitably leaving many unbound copies of dCas9-TET free to affect the genome in a TET-dependent – rather than dCas9-dependent – binding manner. Indeed, our data suggest that dCas9-TET demethylates the *Il33-*002 promoter with a scrambled, non-targeting guide (gRNAscr) (Fig. 1 E-G). This is indicative of a potential ubiquitous and dCas9-independent activity of the fused, over-expressed TET domain in a behavior similar to the demonstrated global methylation by DNMT3A in dCas9-methyltransferase fusions [38]. Thus, whenever a flexibly tethered enzyme is employed for epigenetic editing, it will be difficult to dissociate effects of targeted and nontargeted DNA demethylation on transcription activity.

Finally, the demethylation that is observed with dCas9-TET fusions might be secondary to transcription activation. When we combined our three targeting gRNAs with the well-characterized dCas9-VP64 fusion, VP64 is a potent transcriptional activator originating from the herpes simplex virus [41], we observed broad demethylation of the *Il33-002* promoter (Supp. Fig. S1B-D). This phenomenon suggest that DNA demethylation can in particular instances be secondary to transcription factor recruitment and transcriptional activation (Fig. S1E.) as has been previously reported [92, 93].

Lastly, there are examples in which dCas9 or another targeting protein either bears a catalytically inactive form of TET or the domain is altogether missing, and mild demethylation is still observed [19, 20, 30, 101]. We propose that, in some cases, this demethylation stems from the lingering transactivation capacity of the mutated TET domain (discussed above) followed by demethylation as a consequence of activation, such as the demethylation caused by VP64 activation (Fig. S1E). Alternatively, as we demonstrate here (Fig. 2) binding of dCas9 blocks DNA methyltransferase catalyzed methylation. This therefore obscures the true contribution of TET proteins to demethylation. It is in fact possible that most of the demethylation triggered by dCas9-TET fusions seen in dividing cells stems from the simple steric interference with DNA methyltransferase activity as we demonstrate in this study.

In sum, in the study of causality, the manipulation of a lone variable is a fundamental requirement. The array of transcriptional consequences introduced by TET proteins renders them unable to isolate 5-methylcytosine demethylation as the lone variable and therefore the issue of the causal and functional role of site-specific demethylation in transcriptional activation remains unresolved.

We instead propose and demonstrate here a previously unrecognized capacity of dCas9 to prevent DNA methylation with high efficacy at fairly small, precise regions and, more importantly, free from any fused eukaryotic enzyme that may act independently of the dCas9:gRNA binding activity. Since inhibition of DNA methylation is dependent on tight binding of dCas9 which is also dependent on gRNA target and quality, the risk for nontargeted demethylation is low. Indeed, we did not detect off target demethylation, in contrast to the dCas9-TET fusion construct. We first show that this approach can be implemented as a novel method to map the individual methylated CpGs within a regulatory region which silence transcription using an *in vitro* methylation promoter-reporter transient-transfection assay. Our results demonstrate that three CpG sites within 22 bp of the TSS are sufficient to silence the *Il33-002* promoter while other CpG sites do not contribute to methylation dependent silencing of promoter activity.

We further show that this approach can be applied to trigger site-specific demethylation in dividing cells and that it can be optimized for near-complete removal of DNA methylation from sites that had previously been fully methylated, without perturbing the methylation states of adjacent CpGs in the same promoter to any substantial degree. Though our off-target analyses could be augmented by more comprehensive analyses (whole-genome bisulfite sequencing), our initial observations suggest no off-target demethylation by this approach. Thus, this method could interrogate for the first time the causal role of DNA methylation in silencing gene expression.

We used our method of demethylation to define the role of TSS and proximal promoter methylation of the *Il33-002* gene in its cogent genomic context. We found that demethylation of the *Il33-002* TSS produces a small but significant increase in its expression. Our results confirm what was observed in the transient transfection assay: CpG sites 9 to 11 at the TSS suppress promoter activity. However, dCas9-TET induced 25-fold higher *Il33* expression compared to dCas9 alone when targeted to the same promoter, even though it caused significantly lower demethylation than dCas9 (Fig. 5H). There are several possible explanations for this discrepancy between the fold induction achieved by demethylation and by TET recruitment. First, the fusion of TET to dCas9 is flexible and may allow access to DNA in a wider region, perhaps inducing demethylation in other regulatory regions that are required for more robust expression (Fig. 1 E-G). However, treating cells that have been demethylated at the *Il33-002* TSS CpG sites 9-11 with 5-2’ deoxy-azacytidine doesn’t further induce the gene, while cells that were methylated at 9-11 sites are induced to a level similar to the levels achieved by dCas9. This suggests that the main regulation by DNA methylation occurs at CpGs 9-11 but that the gene is further induced by DNA methylation independent mechanisms that are partially triggered by TET. This illustrates that the results of TET targeting could not be automatically understood as being driven by demethylation and highlights the need for enzyme independent targeted DNA demethylation for understanding the role of DNA methylation.

We then determined whether demethylation of the TSS poises the *Il33-002* promoter for induction by known inducers of this gene. poly(I:C) induces *Il33-002* 300-fold without detectable DNA demethylation and does not induce the demethylated *Il33-002* to a higher level than the methylated *Il33-002* promoter. Thus, induction of *Il33-002* expression is independent of DNA demethylation in the basal promoter. It is possible however that poly(I:C) triggers demethylation in a remote enhancer that wasn’t examined in our study. In contrast, induction by LPS is higher when the basal promoter is demethylated, however LPS induces the promoter whether it is methylated or not suggesting an additive but nonessential effect of demethylation of TSS for LPS induction. What is the role of *Il33-002* promoter methylation? The data is consistent with the idea that this gene is mainly regulated by extracellular signals irrespective of DNA methylation. DNA methylation only silences the residual basal activity of the promoter, perhaps to prevent leaky expression and transcriptional noise in the absence of the appropriate signal. This is consistent with the observation that the ectopically transfected promoter is silenced by DNA methylation (Fig. 2). Therefore, either targeted demethylation or 5-aza-2’-deoxycytidine achieve only a small elevation in expression.

A different paradigm is represented by the *SERPINB5* promoter. Demethylation of the basal promoter on its own has no effect on expression, which remains low (Fig 6 E-F) even when 6 CpGs in the proximal promoter region become completely demethylated (Fig. 6 B-E). However, global demethylation by 5-aza-2’-deoxycytidine induces the activity of this demethylated promoter further than the naturally methylated promoter in control MDA-MB-231 cells, suggesting that expression of this gene is regulated by methylation in the promoter region as well as other regions in *cis* or *trans*. Demethylation of the proximal promoter on its own is insufficient to induce transcription. A possible explanation is that activity of this gene requires activation of a transcription factor that is silenced in these cells and induced by demethylation as we have recently shown [102]. The tumor necrosis factor (*Tnf*) gene exhibits a proximal TSS promoter region that is highly methylated in NIH-3T3 cells. The gene is highly induced and its proximal promoter region is demethylated by 5-aza-2’-deoxycytidine (Fig. 6G). In contrast to the large induction of expression of by 5-aza-2’-deoxycytidine, demethylation of 10 CGs proximal to the TSS (Fig. S9) using the targeted dCas9 method did not turn on the gene (Fig. 6H, grey bars). Here, as was the case with the *SERPINB5* promoter, 5-aza-2’-deoxycytidine treatment of cells bearing dCas9-demethylated *Tnf* TSS region resulted in higher induction of expression than treated control cells bearing a methylated *Tnf* TSS (Fig. 6H). These experiments illustrate the importance of studying demethylation of specific sites *per se* to truly understand their contribution to gene expression control.

Finally, in a manner dissimilar to the other genes examined in this study, apparent demethylation of the large, highly-methylated *FMR1* repeat region in Fragile X syndrome patient fibroblasts did induce basal transcription of the *FMR1* gene up to a 110-fold in one cell pool suggesting that, in this case, methylation of the repeat element plays a large role in silencing of the gene.

In summary, we developed a tool that allows site-specific demethylation of a narrow region of DNA by physical blocking of DNMTs without using confounding epigenetic enzymatic activities. This tool allows us for the first time to examine causal relationships between demethylation of specific sites and gene expression in genes at their native positions in the chromatin. Comparing the results obtained using this tool and results obtained using general DNA methylation inhibitors reveals that the role of DNA methylation at specific sites might have been previously overestimated by confounded techniques. Our study demonstrates the need for the careful causational investigation of the role of DNA methylation of different regions *per se* by an unconfounded tool. We hope that this tool can be used to attribute causality to DNA methylation changes not only in fundamental physiological gene transcription, but also under different specific physiological and pathological conditions mediated by changes in extracellular signals and changes in the milieu of cellular transcription factors in order to begin to reveal the true extent, the nature, and the diverse contribution of DNA methylation at different regions to gene regulation.

## Methods

### gRNA design and synthesis

To maximize likelihood of on-target efficiency and minimize off-target binding, gRNAs were designed using three online tools with distinct scoring algorithms: Off-Spotter, CCTOP, and CRISPR Design [50, 103, 104]. Final gRNAs were chosen based on highest cumulative rank and location in the promoter. The scrambled gRNA sequence was obtained from pCas-Scramble (Origene). For *in vitro* assays, gRNAs were *in vitro* transcribed with the GeneArt^TM^ Precision gRNA Synthesis Kit (Thermo Fisher Scientific) according to manufacturer protocol and using primers listed in **Supplementary Table S2.** Due to a lack of available kit compatible with S. aureus gRNAs, SA-gRNA1-4 and SP-gRNA1-4 were generated by a custom T7 *in vitro* transcription protocol (dx.doi.org/10.17504/protocols.io.dwr7d5) modified to replace the S. pyogenes scaffold sequence with that of S. aureus. (primers in **Supplementary Table S3**). Lentiviral gRNAs were first produced according to the protocol by Prashant Mali [105]. Briefly, 455bp double stranded DNAs containing the human U6 promoter, gRNA sequence, gRNA scaffold, and termination signal were ordered as gBlock Gene Fragments (IDT). These were re-suspended, amplified with Taq Polymerase (Thermo Fisher Scientific) according to manufacturer protocol and using primers listed in **Supplementary Table S2**, extracted from an agarose gel with the QIAEX II Gel Extraction Kit (QIAGEN), and inserted into pCR®2.1-TOPO (Thermo Fisher Scientific) for 30 minutes at room temperature. A lentiviral backbone was obtained from Addgene (pLenti-puro, Addgene #39481) and the CMV promoter was removed to prevent aberrant transcription by digesting the plasmid with ClaI-HF and BamHI-HF (NEB), gel extracting, removing DNA overhangs with the Quick Blunting™ Kit (NEB), and circularization with T4 Ligase (Thermo Fisher Scientific) for 1 hour at 22°C. The resulting promoterless pLenti-puro plasmid was then digested with EcoRI and the 5’ phosphates were removed with Calf Intestinal Alkaline Phosphatase (Thermo Fisher Scientific) to facilitate efficient ligation of the EcoRI-flanked gRNA scaffold. Resulting clones were Sanger sequenced with pBABE 3’ sequencing primer to ensure proper gRNA sequence and orientation (Génome Québec). The gRNAs targeting *SERPINB5* and *Tnf* were created by site-directed mutagenesis of pLenti-Il33gRNA6-puro using primers listed in **Supplementary Table S2** and the Q5® Site-Directed Mutagenesis Kit (NEB) according to manufacturer protocol. The *HNF4A*-targeting gRNA is from the genome-scale CRISPR knock-out (GeCKO) v2 library [106] (purchased as lentiviral plasmid from Genscript) and the FMR1-targeting gRNA from the Jaenisch lab was obtained from Addgene (pgRNA-CGG, Addgene #108248).

### Site-specific *in vitro* DNA methylation

First, a dCas9:gRNA ribonucleoprotein complex was formed with the following mixture: 14 µL nuclease-free water, 3 µL Cas9 Reaction Buffer (Applied Biological Materials Inc.), 7.5 µL 300 nM CpG-targeting *in vitro* transcribed gRNA or non-CpG-targeting control gRNA, and dCas9 recombinant protein (Applied Biological Materials Inc.). After 10 minutes at room temperature, 3 µL of 30 nM Il33-pCpGl was added to the reaction, which was then transferred to 37°C to allow dCas9:gRNA complex binding to DNA. After 1 hour, the following mixture was added to the reaction: 145 µL nuclease-free water, 17 µL NEBuffer™ 2, 5 µL 32mM S-Adenosyl methionine (NEB) (final concentration 0.8 mM), and 3 µL (12 units) M.SssI methyltransferase (NEB). This solution was pre-warmed to 37°C before addition to prevent interference with dCas9:gRNA binding to the DNA. After 4 hours of incubation at 37°C, 1 µL of 20 mg/mL Proteinase K (Roche) was added and the temperature was raised to 64°C for an additional 4 hours.

### DNA Isolation, bisulfite conversion, bisulfite-cloning, and pyrosequencing

Plasmid DNA was then recovered by phenol-chloroform extraction and precipitation in ethanol overnight. DNA was washed one time with 70% ethanol, dried, and re-suspended in 30 µL nuclease-free water. Genomic DNA was extracted from cells by resuspension in 400 µL DNA lysis buffer (100mM Tris, pH 7.5, 150 mM NaCl, 0.5% SDS, 10 mM EDTA), phenol-chloroform extraction, ethanol precipitation and washing as described previously. Following DNA extraction, bisulfite conversion was conducted according to manufacturer protocol with the EZ DNA Methylation-Gold Kit (Zymo Research) using 5 µL of *in vitro* methylated plasmid DNA or 1.5 µg genomic DNA measured with the Qubit dsDNA BR Assay (Thermo Fisher Scientific). 1 µL of bisulfite-converted DNA was amplified with HotStar Taq DNA polymerase (QIAGEN) in a 25 µL reaction using the primers designed with MethPrimer [107] and listed in **Supplementary Table S2**. Pyrosequencing samples were processed in the PyroMark Q24 instrument according to protocols designed by the PyroMark Q24 software (QIAGEN). Sequencing primers were designed with Primer3 [108]. Alternatively, amplicons were cloned into pCR®4-TOPO (Thermo Fisher Scientific) for 30 minutes at room temperature and transformed into TOP10 competent cells (Thermo Fisher Scientific) prior to plasmid isolation with the High-Speed Plasmid Mini Kit (Geneaid) and Sanger sequencing (Eurofins Genomics) using the M13R sequencing primer. All oligonucleotides used in this study were purchased from Integrated DNA Technologies.

### Luciferase Assay

8.0×10^4^ NIH-3T3 cells (*Il33* experiments) or 1.2×10^5^ HEK293 cells (TET co-transfection) were plated in a 6-well plate (Corning) 24 hours prior to transfection. 1 µg (*Il33*) or 100 ng (SV40) plasmid DNA from the *in vitro* methylation reactions were transfected with 3 µL (*Il33)* or 1 µL (SV40) X-tremeGENE 9 transfection reagent (Roche) diluted in 50 µL of Opti-MEM medium (Gibco). Luciferase assays were performed 36 hours after transfection using the Luciferase Reporter Gene Assay, high sensitivity (Roche). Briefly, cells were washed with 1 mL of phosphate-buffered saline (Wisent), detached with scrapers (Thermo Fisher Scientific) after the addition of 150 µL lysis buffer, and transferred to 1.5 mL tubes. After a 15-minute incubation at room temperature, the mixtures were centrifuged for 5 seconds at maximum speed and the supernatant transferred to new 1.5 mL tubes. Two 50 µL volumes per condition were supplemented with 50 µL luciferase assay reagent in disposable glass tubes (Thermo Fisher Scientific) and light emission was measured immediately in the Monolight 3010 luminometer (Analytical Luminescence Laboratory). Sample protein concentration was determined by Bradford Protein Assay (Bio-Rad) and A595 readings were measured in a DU 730 UV-Vis Spectrophotometer (Beckman Coulter). Concentration was determined by comparing to a bovine serum albumin standard curve and luciferase activity was normalized to concentration.

### Plasmids

The original dCas9 plasmid lacking loxP sites was obtained as a dCas9-VP64 fusion (lenti dCAS-VP64_Blast, Addgene #61425). The VP64 domain was removed by digestion with BamHI-HF and BsrGI-HF, blunting with the Quick Blunting™ Kit (NEB), and circularization with T4 Ligase (Thermo Fisher Scientific) for 1 hour at 22°C. Following transformation, plasmids were isolated from ampicillin-resistant clones (High-Speed Plasmid Mini Kit, Geneaid) and Sanger sequenced to identify plasmids that maintained the blasticidin resistance gene in-frame with dCas9. Floxed dCas9 was purchased as a ready plasmid (pLV hUbC-dCas9-T2A-GFP, Addgene #53191) and primers were designed to amplify a fragment of approximately 500 base pairs when dCas9 is removed with Cre recombinase (**Supplementary Table S2**). The Cre-containing plasmid was obtained from Addgene (pLM-CMV-R-Cre, Addgene #27546). A fragment encoding the CMV promoter and mCherry-T2A-Cre-WPRE was excised by NdeI and SacII (Thermo Fisher Scientific) and transferred to the pLenti6/V5-DEST™ Gateway™ Vector (Thermo Fisher Scientific) bearing a blasticidin resistance cassette (Thermo Fisher Scientific) to facilitate antibiotic selection. Lentiviral Fuw-dCas9-Tet1CD-P2A-BFP and Fuw-dCas9-dead Tet1CD-P2A-BFP were obtained from Addgene (Addgene #108245,#108246). Catalytically active Cas9 lentiviral vector was obtained from Genscript as pLentiCas9-Blast. pcDNA3-TET2 (Fig 1J) was generated by amplification of TET2 from human cDNA, TOPO-TA cloning and sequence validation by Sanger sequencing, followed by digestion and ligation into pcDNA3.1 (Thermo Fisher Scientific) using the restriction enzymes XhoI and ApaI. SV40-pCpGl (Fig 1J) was generated by amplification of the SV40 promoter and enhancer region from lenti dCAS-VP64_Blast using primers that added a 5’ BamHI site and a 3’ HindIII site, which were then used for transfer into pCpGl [45] following sequence verification.

### Cell Culture

HEK293T and NIH-3T3 cells (ATCC) were thawed and maintained in DMEM medium (Gibco) supplemented with 10% Premium Fetal Bovine Serum (Wisent) and 1X Penicillin-Streptomycin-Glutamine (Gibco). Cells were grown in a humidified incubator of 5% carbon dioxide at 37°C and cultured in 100mmx20mm tissue culture dishes (Corning) and harvested or passaged by trypsinization (Gibco) upon reaching 80-90% confluency. Clones were isolated by limiting dilution and trypsinization with the aid of cloning rings. Fragile X syndrome fibroblasts (GM05848, Coriell Institute) were maintained as above. Flow cytometry to isolate dCas9-TET/dCas9-deadTET (BFP) and dCas9 (GFP) when antibiotic selection was not an option was performed by Julien Leconte of the Flow Cytometry Core Facility at McGill University Life Sciences complex.

### Lentiviral Production

HEK293T cells were plated at a density of 3.8 x 10^6^ per 100mm dish 24 hours prior to transfection. Cells were transfected using X-tremeGENE 9 transfection reagent (Roche). Briefly, individual lentiviral transfer plasmids were mixed with a packaging plasmid (pMDLg/pRRE, Addgene #12251), envelope protein plasmid (pMD2.G, Addgene #12259), REV-expressing plasmid (pRSV-Rev, Addgene #12253), and the transfection reagent in Opti-MEM medium (Gibco). The mixture was incubated for 30 minutes at room temperature and added in a drop-wise manner to HEK293T cells in 8 mL of fresh DMEM medium in a 100mm dish. Lentiviral particles were harvested by filtering the supernatant through a 0.45 μm disk filter 72 hours after transfection and either used immediately or stored at −80°C. 5 µg/mL Blasticidin S HCl and 1 µg/mL Puromycin Dihydrochloride (Gibco) were used to select for stable transformants.

### RT-qPCR

RNA was isolated from approximately 80% confluent 100mm dishes with 1 mL of Trizol reagent (Thermo Fisher Scientific) following harvest by trypsinization and washing with phosphate-buffered saline (Wisent). RNA extraction was performed according to Trizol manufacturer protocol. Briefly, 200 mL of chloroform was added to 1 mL of Trizol-RNA mixture. The samples were thoroughly vortexed, incubated at room temperature for 2 minutes, and centrifuged for 15 minutes at 12,000 xg at 4°C. The aqueous phase was transferred to a new 1.5 mL tube prior to the addition of 0.5 mL isopropanol and incubation at room temperature for 10 minutes. The samples were centrifuged for 10 minutes at 12,000 xg at 4°C, and washed twice with 75% ethanol, discarding the supernatant each time. The pellets were air dried for 10 minutes and re-suspended in 50 µL DEPC-treated water (Ambion). Concentrations were measured with the Qubit RNA BR Assay (Thermo Fisher Scientific) and 1 µg RNA was used for each reverse transcriptase reaction using M-MuLV Reverse Transcriptase (NEB) according to manufacturer protocol. cDNA was diluted 1:2 (20 µL reverse transcription reaction to 40 µL water) and 2 µL of diluted cDNA was amplified in the LightCycler ® 480 Instrument II (Roche) in a 20 µL reaction containing 10 µL LightCycler ® 480 SYBR Green I Master Mix (Roche) and 0.8 µL each of 10 µM forward and reverse primer listed in **Supplementary Table S2**. Quantification was performed by Roche Lightcycler Software.

### Drug Treatment

5-aza-2’-deoxycytidine (Sigma A3656) was dissolved to 10 mM in sterile water and frozen in one-time-use aliquots at −80°C. Trichostatin A (TSA, Sigma T8552) was dissolved to 1 mM in dimethyl sulfoxide (DMSO, Sigma D8418) and frozen in one-time-use aliquots at −80°C. Lipopolysaccharides from Escherichia coli O55:B5 (Sigma L6529) were diluted to 1mM in phosphate-buffered saline. 5-aza-2’-deoxycytidine and TSA treatment regiment involved 3 treatments every other day with media replacement (5 days total) at specified concentrations and sample collection on the sixth day.

### Off-Target Prediction

Potential off-target sites of *Il33* gRNA3 in the mouse genome were predicted using Cas-OFFinder [109], a program that allows bulges in the RNA and DNA (which Cas9 is known to tolerate) to increase the number of possible off-target sites. Because we were interested in changes in methylation, results were filtered for the presence of a CG at a maximum of 10bp from either end of the gRNA sequence. Of 15 results, 2 differed by 3 mismatches, 9 by 4 mismatches, and 4 by 2 mismatches and a bulge. We developed functional pyrosequencing assays for 4 of these sites.

### Hydroxymethylation quantification

DNA isolated from cells by phenol:chloroform isolation and ethanol precipitation was cleaned on Micro Bio-Spin P-6 SSC columns (Bio-Rad) according to manufacturer protocol. 15 mM KRuO4 (Sigma) was prepared by dissolving 0.153g in 50 mL of 0.05M NaOH and thawed freshly for each oxidation reaction. 1 µg cleaned DNA was incubated in a 19 µL volume reaction in a PCR tube with 0.95 µL 1M NaOH at 37 °C in a shaking incubator for 0.5 hr. The sample was cooled immediately in an ice-water bath for 5 min prior to the addition of 1 µL ice-cold 15 mM KRuO4 and incubation in an ice-water bath for 1 hr with vortexing every 20 min. A second oxidation was performed by the addition of 4 µL 0.05 M NaOH, incubation at 37 °C in a shaking incubator for 0.5 hr, following by cooling, addition of 1 µL ice-cold 15 mM KRuO4, and incubation in ice-water bath with occasional vortexing as before. Oxidized DNA was cleaned again on Micro Bio-Spin P-6 SSC columns and the DNA was subjected to bisulfite conversion and pyrosequencing. Control reactions were done in parallel in which 15 mM KRuO4 was replaced by 0.05 M NaOH and percent hydroxymethylation was quantified as the decrease in methylated fraction in oxidized DNA as compared to control DNA.

### Chromatin Immunoprecipitation

150mm tissue culture dishes containing 90% confluent NIH-3T3 cells from each experimental condition were cross-linked by the direct addition of formaldehyde to a 1% final concentration. The dishes were incubated for 10 minutes at room temperature with constant agitation. The reaction was quenched by the addition of glycine to a final concentration of 0.125 M and incubated for an additional 5 minutes at room temperature with constant agitation. Cross-linking solution was aspirated and cross-linked cells were washed three times with 10 mL ice-cold phosphate-buffered saline (PBS). 10 mL of ice-cold PBS was added and cells were scraped into suspension by a rubber cell scraper. Cross-linked cells were pelleted at 800xg at 4°C in 15mL falcon tubes, the supernatant removed, and the cells were lysed in 300 µL ice-cold lysis buffer (1% SDS, 10 mM EDTA, 50 mM Tris), pipeted thoroughly, incubated for 15 minutes on ice, and immediately sonicated on the Bioruptor (Diagenode) in 1.5 mL Eppendorf tubes at high power for three 10-minute cycles of 30 seconds on and 30 seconds off, replacing warmed water with ice-cold water and minimal ice between each cycle. Sonicated samples were centrifuged at 4°C for 16,000xg for 5 minutes, and supernatant was transferred to a clean 1.5 mL Eppendorf tube, with 30 µL set aside for shearing efficiency analysis. The remaining supernatant was diluted with a 9X volume of dilution buffer (16.7mM TrisHCl pH 8.0, 1.2mM EDTA, 167mM NaCl, 0.01% SDS, 1.1% Triton X-100) and precleared with washed Dynabeads Protein G (Thermo Fisher) for 2 hrs at 4°C on a nutator. Using a magnetic rack, 1% of pre-cleared chromatin was set aside for input and 5 µg Monoclonal ANTI-FLAG® M2 antibody (to capture FLAG-tagged dCas9) or IgG (Sigma) was added to the remaining volume and then incubated at 4°C on a nutator overnight. 60 µL of washed (3X with Tris-EDTA – 10mM Tris pH=8, 1mM EDTA – and 3X with RIPA – 20 mM Tris, 2 mM EDTA, 150 mM NaCl, 1% Triton X, 0.1% SDS, 0.5% deoxycholate) Dynabeads were added to each sample and incubated at 4°C on a nutator for 4 hrs. Beads were then washed with 1mL each as follows: 2X with low salt wash buffer (0.1% SDS, 1% Triton-X, 2mM EDTA, 20mM Tris, 150mM NaCl), 2X with high salt wash buffer (same as low except 500 mM NaCl), 2X with LiCl wash buffer (0.25M LiCl, 1% NP-40, 1% deoxycholate, 1mM EDTA, 10mM Tris, pH 8.0), and 2X with Tris-EDTA. All buffers contained 1X cOmplete™ Protease Inhibitor Cocktail (Sigma). DNA was eluted by the addition of 100 µL elution buffer (1% SDS, 0.1M NaHCO3), vortexing vigorously, and 15-minute incubation at room temperature with constant agitation before transferring to a clean 1.5 mL tube. This was repeated twice for a final volume of 200 µL and the input fraction was adjusted to the same volume with elution buffer. Reverse cross-linking (0.2M final concentration of NaCl, 65 °C overnight) was performed for all samples, followed by standard treatment with RNAse A, proteinase K, and phenol:chloroform cleanup followed by ethanol precipitation. Clean DNA was then quantified by qPCR and enrichment in the immunoprecipitated samples was calculated as fraction of input. Nonspecific (IgG) antibody and qPCR primers of unbound regions were used as controls for effective immunoprecipitation.

## Acknowledgements

This study was funded by the Canadian Institutes of Health Research (PJT159583). D.M.S was supported by fellowships from the McGill University Faculty of Medicine (Friends of McGill Fellowship; JP Collip Fellowship in Medical Research; James Frosst Fellowship).

## Supplementary Figures

**Supplementary Figure S1.**
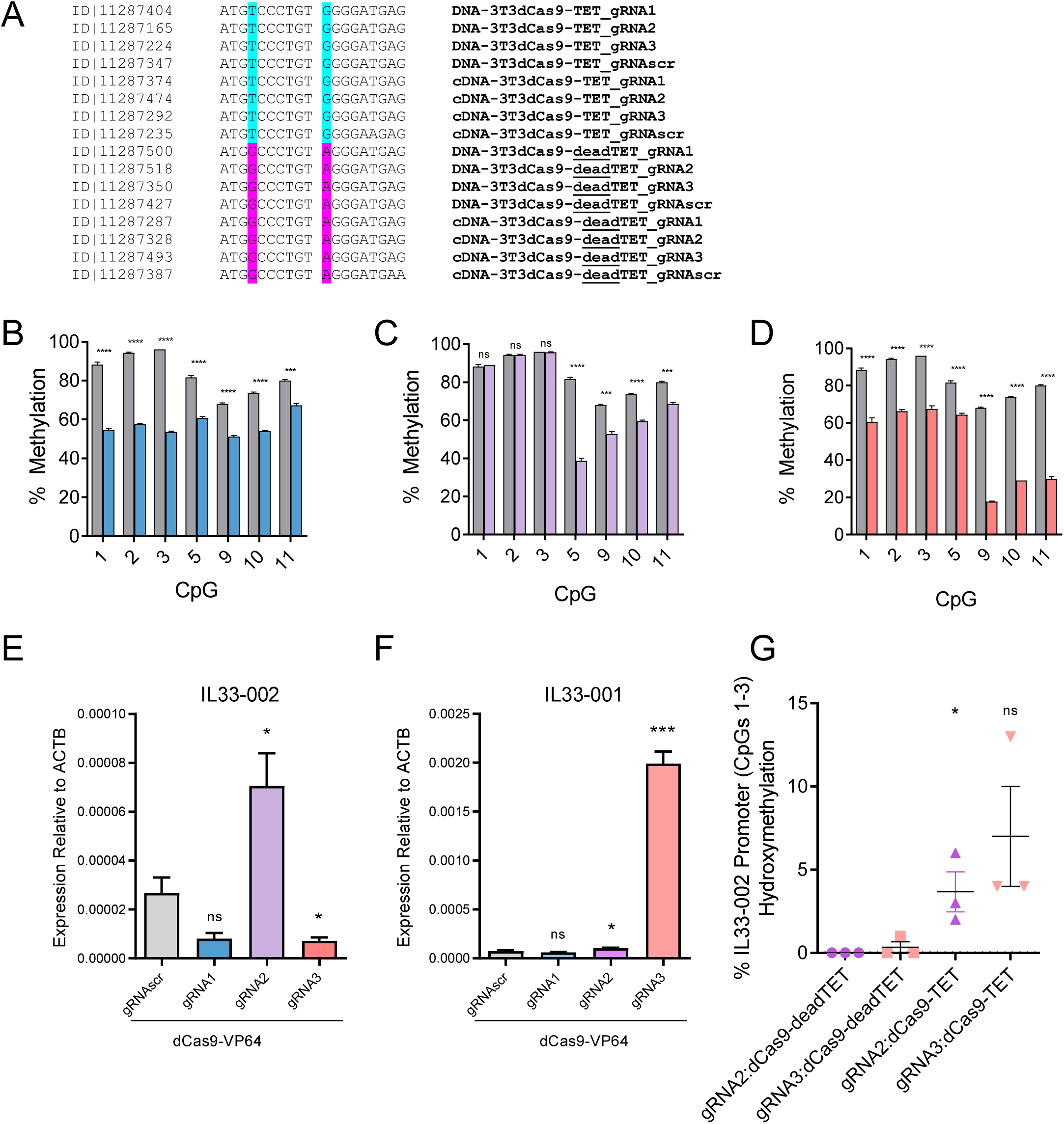
(A) Aligned Sanger sequencing results of region of TET bearing the inactivating mutation in deadTET controls from one representative cell line from triplicate treatments in Fig. 1 with Sanger ID (left), DNA sequence (middle), and source of DNA (right). (B-D) Percent of DNA methylation assayed by bisulfite-pyrosequencing at 7 targeted CpGs in NIH-3T3 cells treated with dCas9-VP64 and either gRNA1 (B; blue), gRNA2 (C; purple), gRNA3 (D; pink) or gRNAscr (grey; identical data in B-D, shown for comparison). E-F. Expression of Il33-002 (E) and Il33-001 (F) quantified by RT-qPCR and normalized to *Actb* expression in NIH-3T3 stably expressing one of 4 gRNAs and dCas9-VP64. G. Percent of DNA hydroxymethylation assayed by KRuO4 oxidation of DNA followed by bisulfite-pyrosequencing in parallel with unoxidized controls and calculated as decrease in methylation after oxidation at CpGs 1, 2, and 3, averaged, which are distant from the gRNA2 (purple) and gRNA3 (pink) binding sites, under the specified stable treatments in NIH-3T3 cells (x-axis). * indicates statistically significant difference of p<0.05, ** of p<0.01, *** of p<0.001, **** of p<0.0001, and ns = not significant (Student’s t-test, with Holm-Sidak correction if number of tests is greater than 3).

**Supplementary Figure S2.**
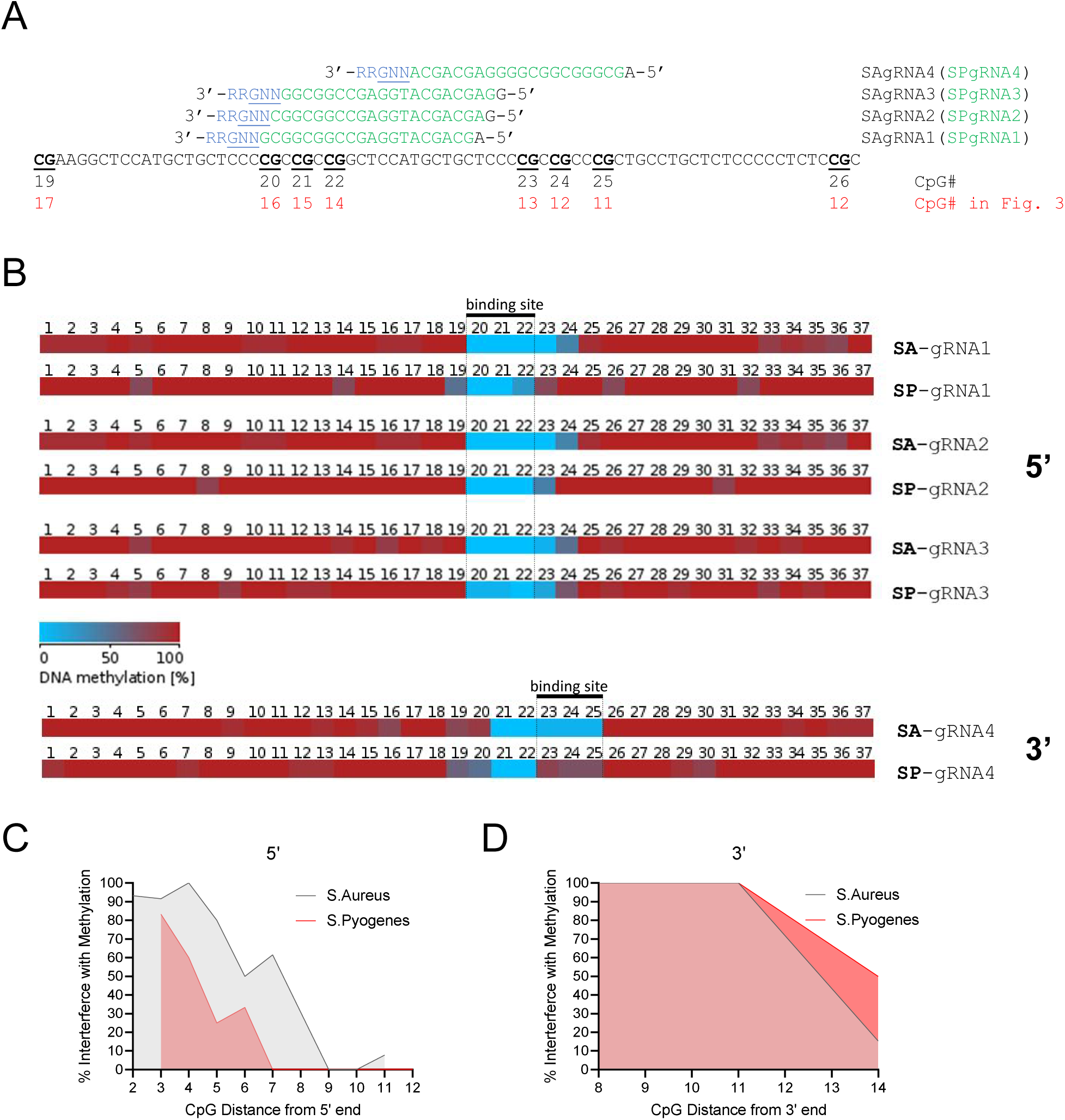
(A) Diagram and sequence of *CDKN2A* region targeted by additional gRNAs. Due to the display of the reverse complement sequence of Fig. 3, CpGs are numbered differently (black) but Fig. 3 numbering system is shown below in red. gRNA sequences are shown in green, where the entire green sequence represents the S. pyogenes gRNA and the addition 5’ nucleotide in black represents the additional nucleotide needed for S. aureus gRNAs. S. aureus PAM site is shown in blue, with the first 3 nucleotides (5’ to 3’) represent the NGG PAM of the S. pyogenes gRNA. (B) Each horizontal row depicts a heatmap of average DNA methylation at each numbered CpG over 10-20 (except SP-gRNA4, 4 clones) individual strands of DNA (bisulfite-converted clones) where light blue represents 0% methylation and dark red represents 100% methylation. The CpGs within the binding site of the labeled gRNA are labeled and enclosed in dashed lines. gRNAs1-3 interrogate DNA methylation interference of 5’ proximal CpGs and gRNA4 interrogates that of 3’ proximal CpGs. Lowly methylated strands of DNA (poor M.SssI methylation) and strands with unaffected binding sites (unbound by dCas9) were excluded from the analysis because efficacy was not under evaluation. (C-D) Data from (B) transformed into a percent methylation as a function of CpG distance in base pairs from the 5’ (C) or 3’ (D) end of the gRNA sequence (including PAM) and S. aureus (grey) or S. pyogenes (pink) across gRNAs 1-3 (C) or gRNA4 (D).

**Supplementary Figure S3.**
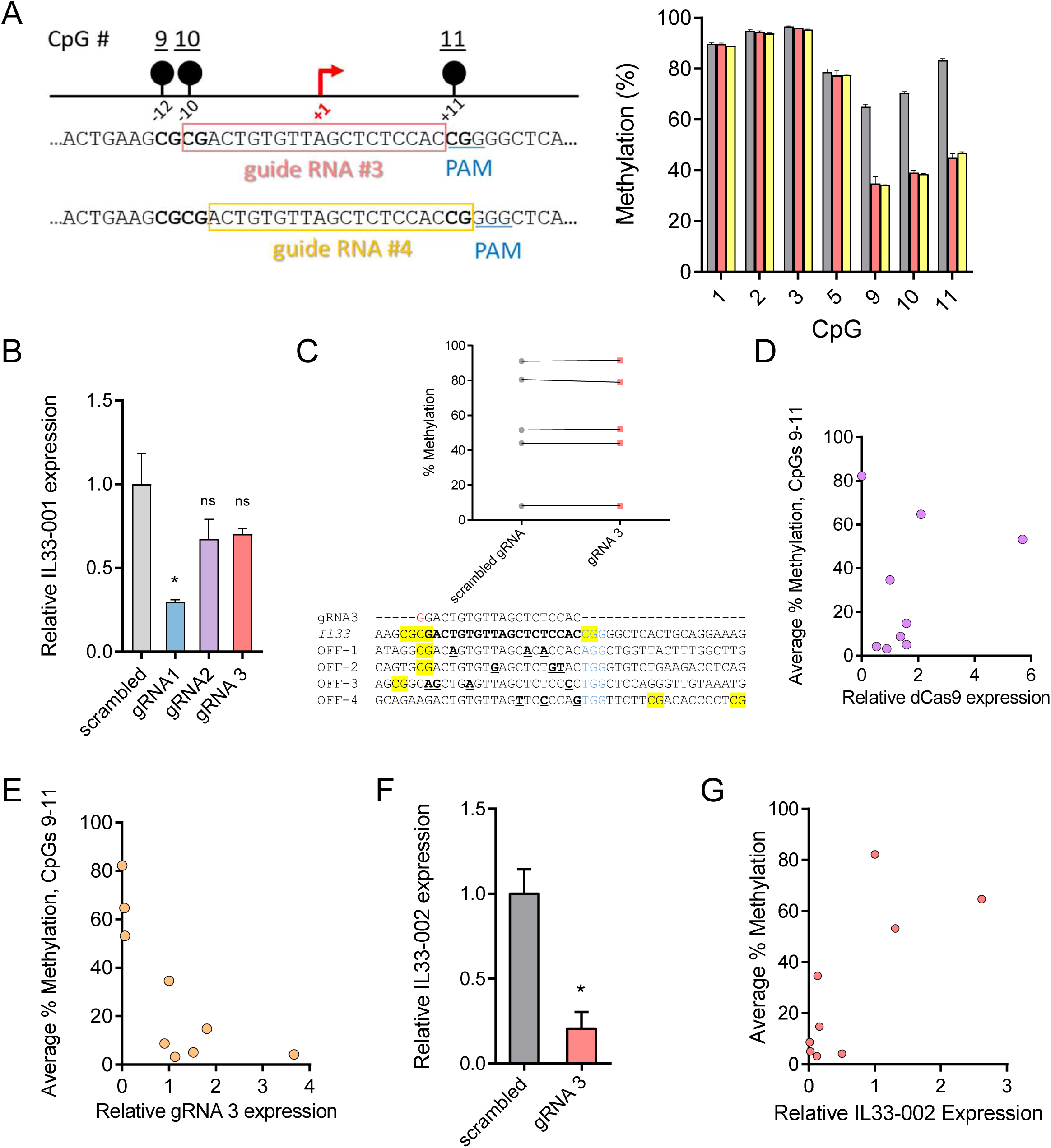
(A) (Left) Diagram of *Il33-002* promoter with location of gRNA3 and gRNA4. TSS is marked by a red arrow and CpGs are marked by black circles. In sequence below, PAM represents protospacer adjacent motif, gRNA sequences are boxed, and CpGs are bolded. (Right) Methylation levels assessed by pyrosequencing of NIH-3T3 cells expressing dCas9 and gRNA3 (pink) or gRNA4 (yellow). Values displayed as mean +/- SEM of 3 biological replicates. (B) *Il33-001* expression in NIH-3T3 cell lines stably expressing gRNAscr or one of 3 *Il33-002*-targeting gRNAs in combination with dCas9, assayed by qRT-PCR and normalized to *Actb* expression (n=3). (C) Comparison of methylation levels, assayed by pyrosequencing, of 5 top off-target CpGs in NIH-3T3 cell lines stably expressing scrambled gRNA or gRNA 3 in combination with dCas9. (D) Correlation of 9 dCas9:gRNA3 clones from (Fig 4E); x-axis displays dCas9 expression normalized to GAPDH expression; y-axis displays average methylation at CpGs 9, 10, and 11 in each clone assayed by pyrosequencing (r=0.1982, p=0.6091). (E) Correlation of 9 dCas9:gRNA3 clones from (Fig 4E); x-axis displays gRNA3 expression normalized to GAPDH expression (F); y-axis displays average methylation at CpGs 9, 10, and 11 in each clone assayed by pyrosequencing (r= −0.7307, p<0.05). (F) *Il33-002* expression in NIH-3T3 cell lines stably expressing scrambled gRNA or gRNA3 in combination with dCas9, assayed by qRT-PCR and normalized to Actb expression (n=3, * indicates p<0.05 vs gRNAscr, t-test). (G) Scatter plot of 9 dCas9:gRNA3 clones from (4E); x-axis displays relative Il33-002 expression as assayed in (F); y-axis displays average methylation at CpGs 9, 10, and 11 in each clone assayed by pyrosequencing (r=0.7395, p=0.0228). * indicates statistically significant difference of p<0.05, ** of p<0.01, *** of p<0.001, **** of p<0.0001, and ns = not significant (Student’s t-test, with Holm-Sidak correction if number of tests is greater than 3).

**Supplementary Figure S4.**
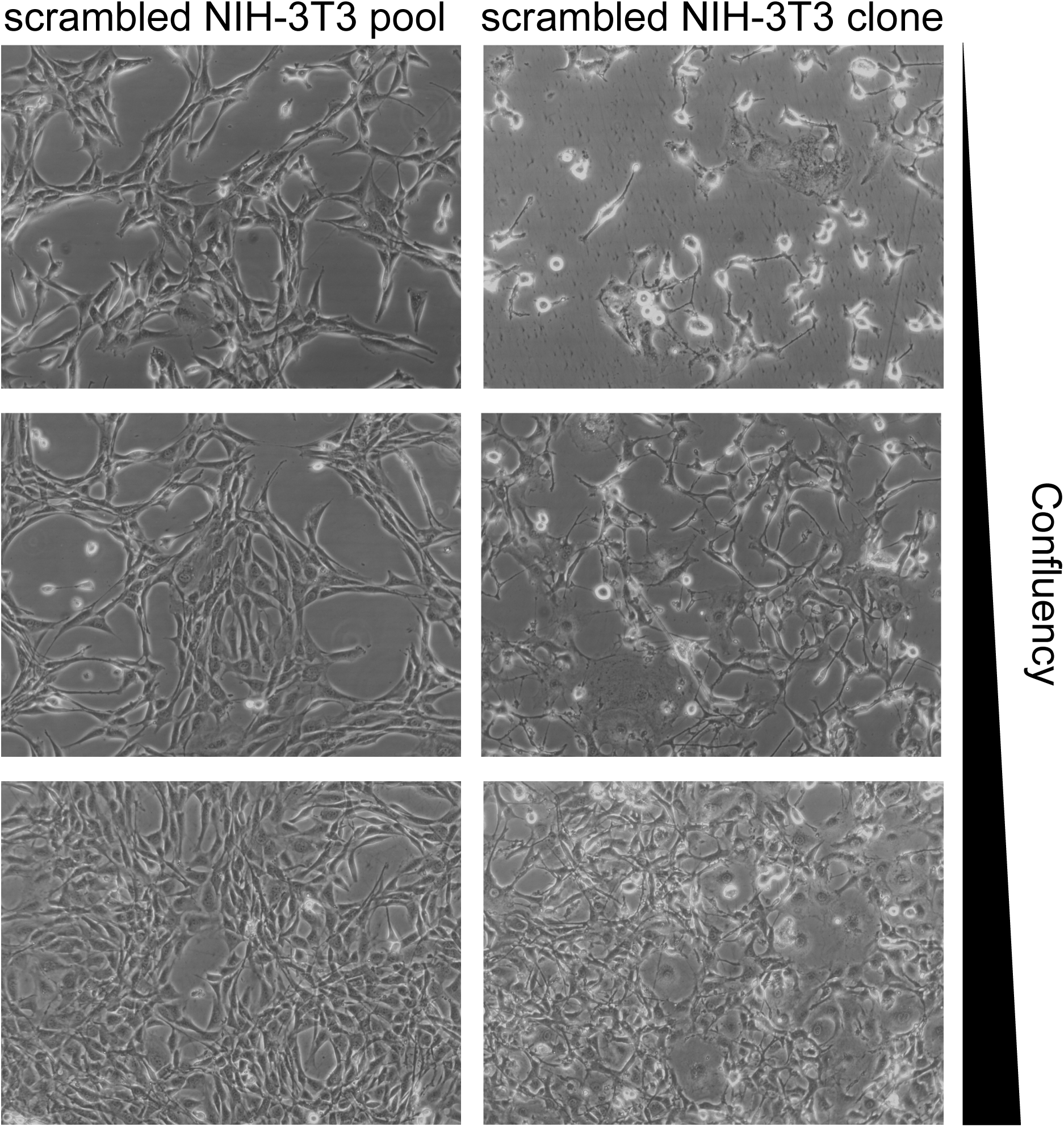
Light microscopy images demonstrating the appearance of normal healthy NIH-3T3 pools (3 left panels) with increasing confluency downwards. In comparison, morphological irregularities can be seen after clonal isolation from the same source cells in 3 distinct clonal populations at 3 different levels of confluency.

**Supplementary Figure S5.**
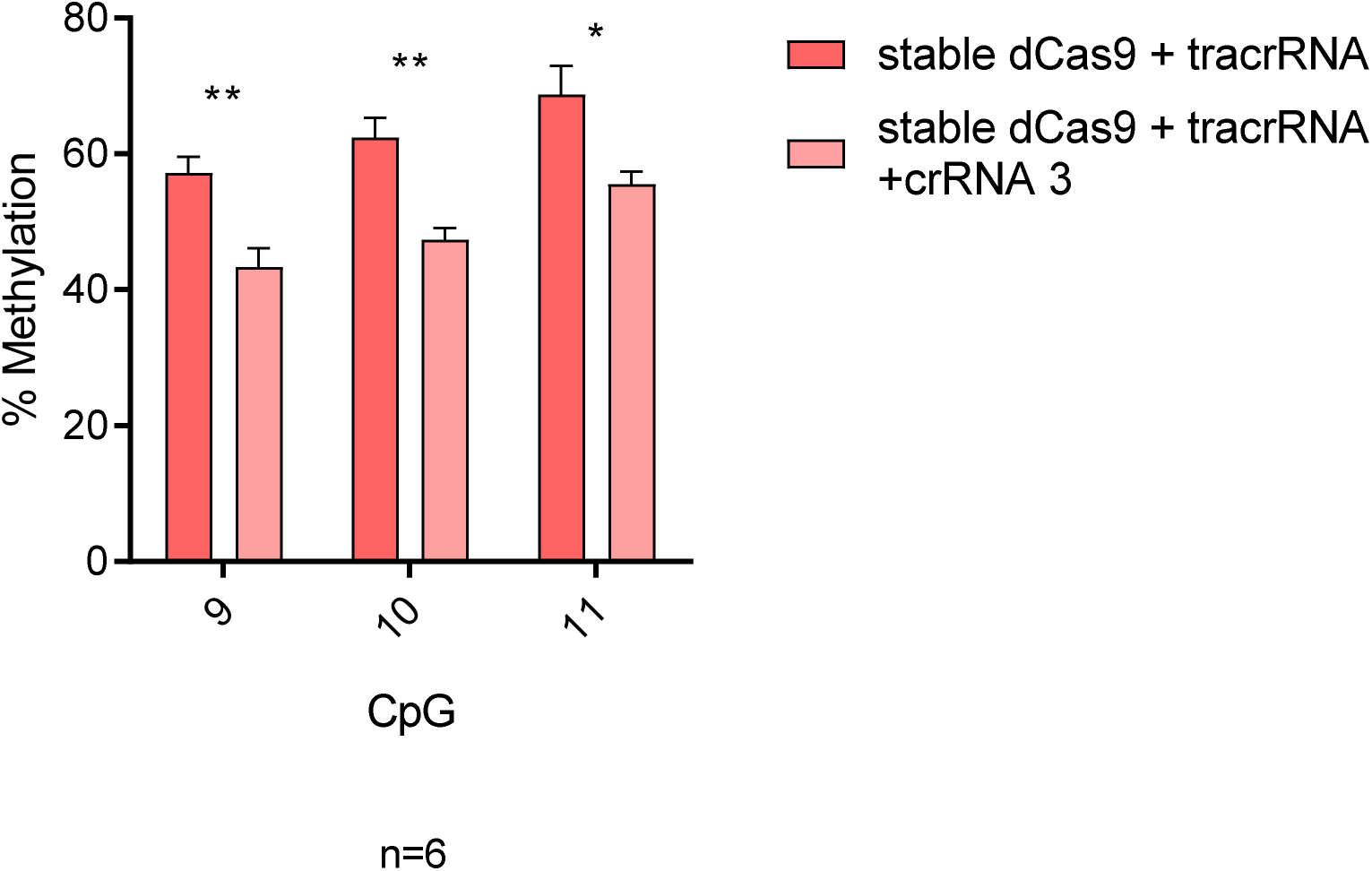
Methylation levels assessed by bisulfite-pyrosequencing of target CpGs 9, 10, and 11 after NIH-3T3 cells stably expressing dCas9 were transiently transfected (Xtremegene siRNA transfection reagent, Sigma) with either tracrRNA alone (red) or tracrRNA and crRNA3 (pink), a two-component version of gRNA3 (n=6). * indicates statistically significant difference of p<0.05, ** of p<0.01, *** of p<0.001, **** of p<0.0001, and ns = not significant (Student’s t-test, with Holm-Sidak correction).

**Supplementary Figure S6.**
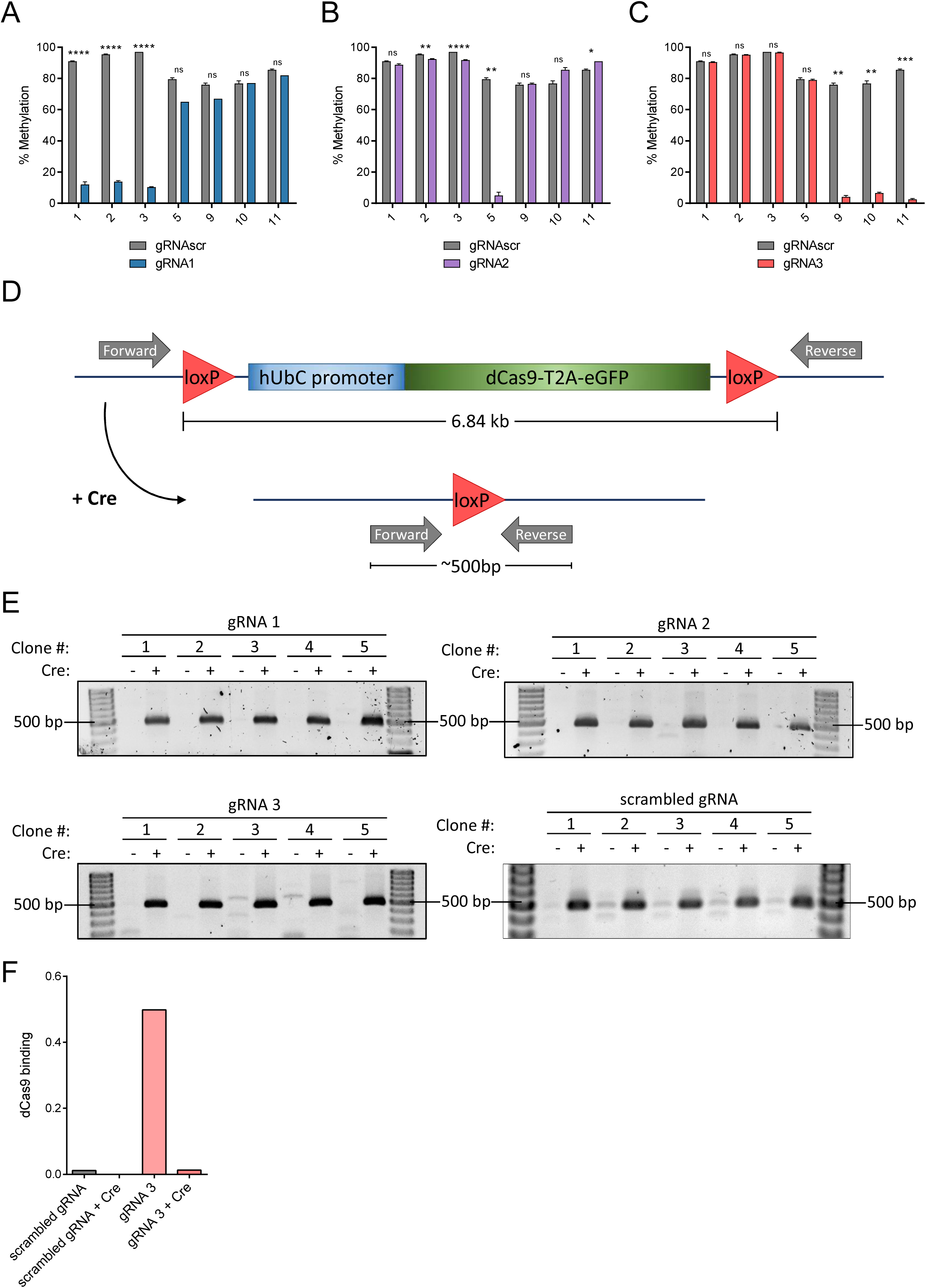
(A-C) DNA methylation analysis by pyrosequencing of NIH-3T3 cells stably expressing gRNA1 (A), gRNA2 (B), or gRNA3 (C) and floxed dCas9 in order to validate targeted demethylation. (D) Diagram of the dCas9 expression construct. It is flanked by loxP sites that facilitate recombination and deletion by Cre recombinase. Forward and reverse PCR primers lie outside the loxP sites such that a 6.84 kb product could be made when Cre is not present in the cells and if PCR extension times are increased to allow this product to form. After removal of the dCas9 expression cassette by Cre recombinase, the same PCR primers create a product of approximately 500 base pairs in size. (E) Agarose gels showing recombination-dependent PCR products using the primers in (D) in cell lines stably expressing dCas9 and each indicated gRNA after treatment by empty virus (-) or Cre recombinase (+). A 500 base pair product is visible in each Cre-containing lane. Primers are listed in Supplementary Table S2. (F) Chromatin immunoprecipitation of cells from Fig. S6E with antibody for dCas9 (anti-Flag) followed by qPCR using primers surrounding the *Il33-002* TSS. dCas9 binding is expressed as percent input (n=1). * indicates statistically significant difference of p<0.05, ** of p<0.01, *** of p<0.001, **** of p<0.0001, and ns = not significant (Student’s t-test, with Holm-Sidak correction if number of tests is greater than 3).

**Supplementary Figure S7.**
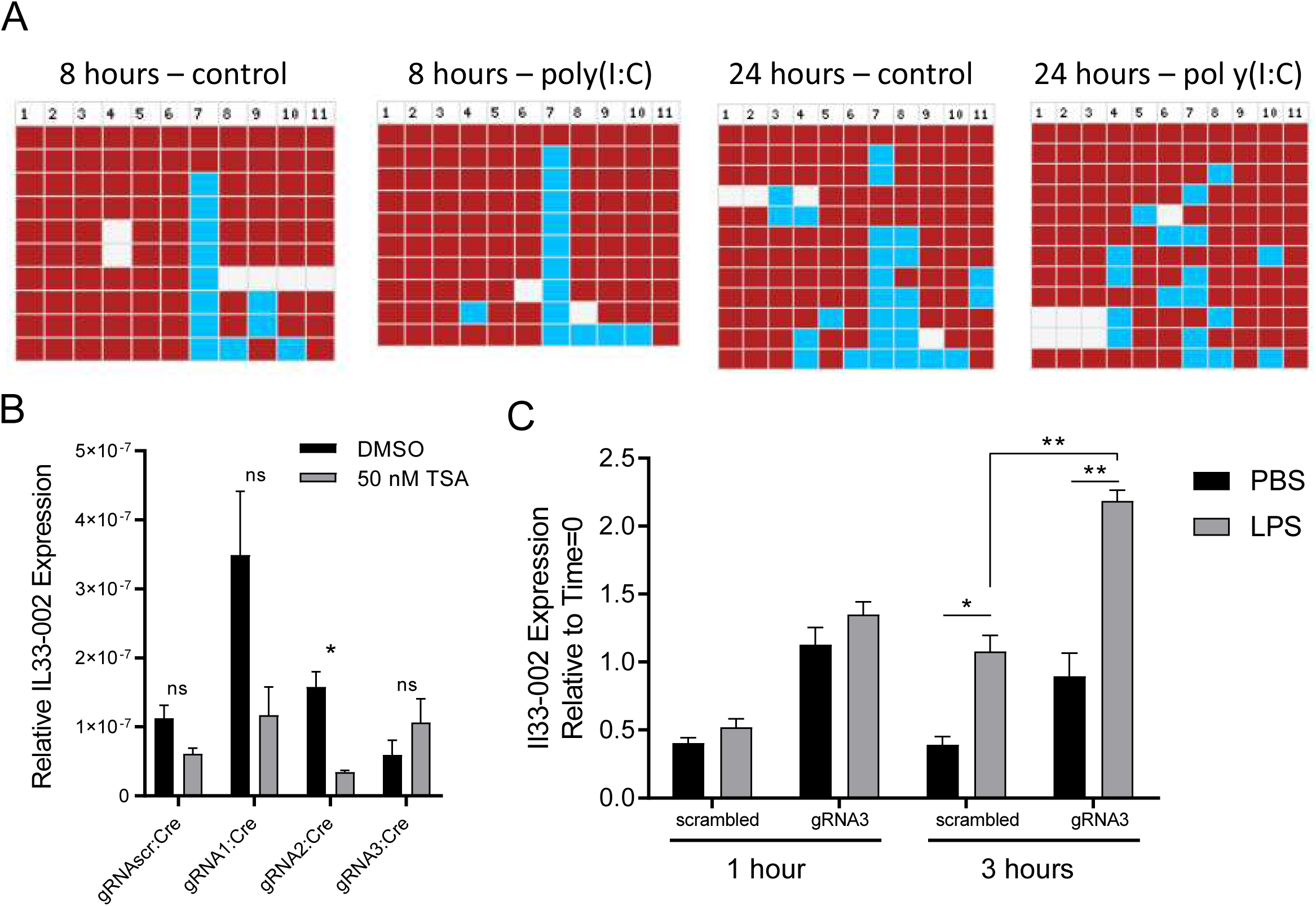
(A) Bisulfite-cloning and sanger sequencing analysis of the *Il33-002* promoter in NIH-3T3 cells treated with 1 µg/mL poly(I:C) or water control for 8 or 24 hours. Each horizontal row is one strand of DNA. Numbers indicate the CpG in the promoter. Red squares indicate methylated CpGs, blue squares indicate unmethylated CpGs, and white squares indicate a lack of data due to sequencing failure. (B) *Il33-002* expression in 50nM TSA or vehicle (DMSO) treated NIH-3T3 cell lines stably expressing gRNAscr, gRNA1, gRNA2, or gRNA3 under high-puromycin conditions in combination with dCas9, followed by dCas9 removal by Cre recombinase as assayed by qRT-PCR and normalized to *Actb* expression (n=3). (C) *Il33-002* expression in 100 ng/mL LPS or vehicle (PBS) treated NIH-3T3 cell lines stably expressing gRNAscr or gRNA3 under high-puromycin conditions in combination with dCas9, followed by dCas9 removal by Cre recombinase, as assayed by qRT-PCR and normalized to *Actb* expression, either 1 or 3 hours after treatment and displayed relative to expression measured at time=0 (n=3). * indicates statistically significant difference of p<0.05, ** of p<0.01, *** of p<0.001, **** of p<0.0001, and ns = not significant (Student’s t-test, with Holm-Sidak correction if number of tests is greater than 3).

**Supplementary Figure S8.**
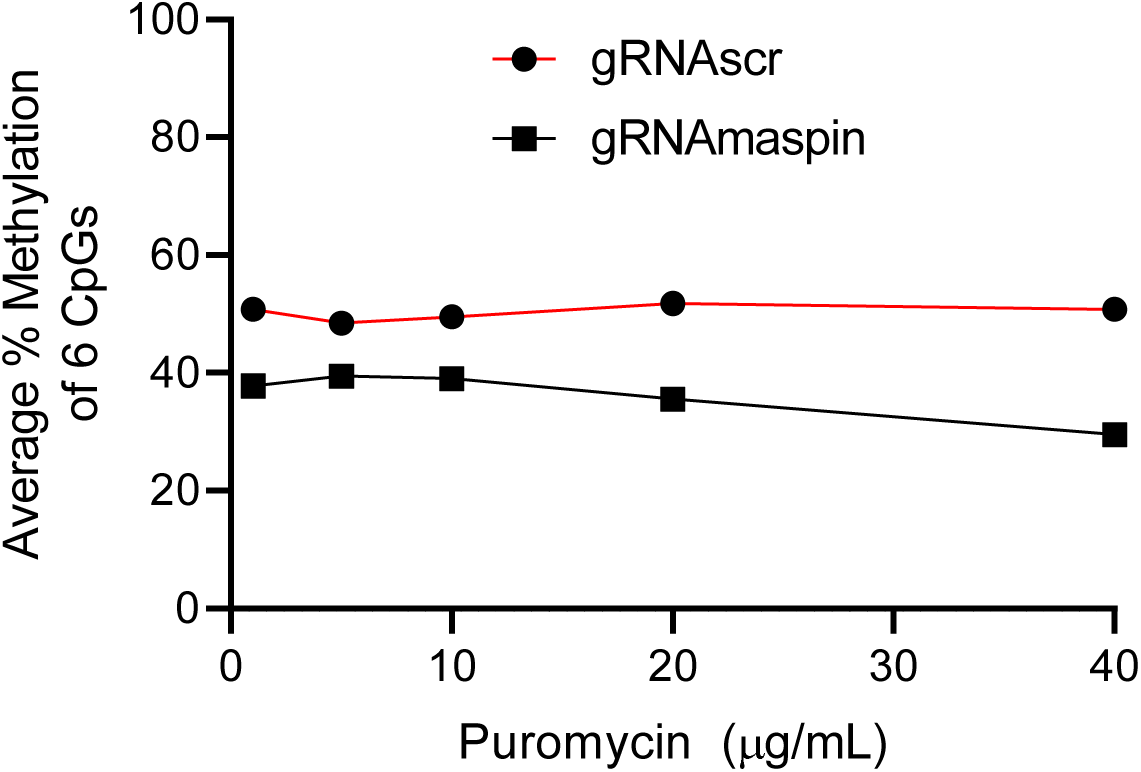
DNA methylation levels assessed by bisulfite-pyrosequencing of CpGs 1-6 in the *SERPINB5* promoter in MDA-MB-231 cell lines stably expressing dCas9 and either gRNAscr (red) or gRNAmaspin (black), averaged across all 6 CpGs and plotted as a function of increasing puromycin concentration (n=1 per puromycin concentration).

**Supplementary Figure S9.**
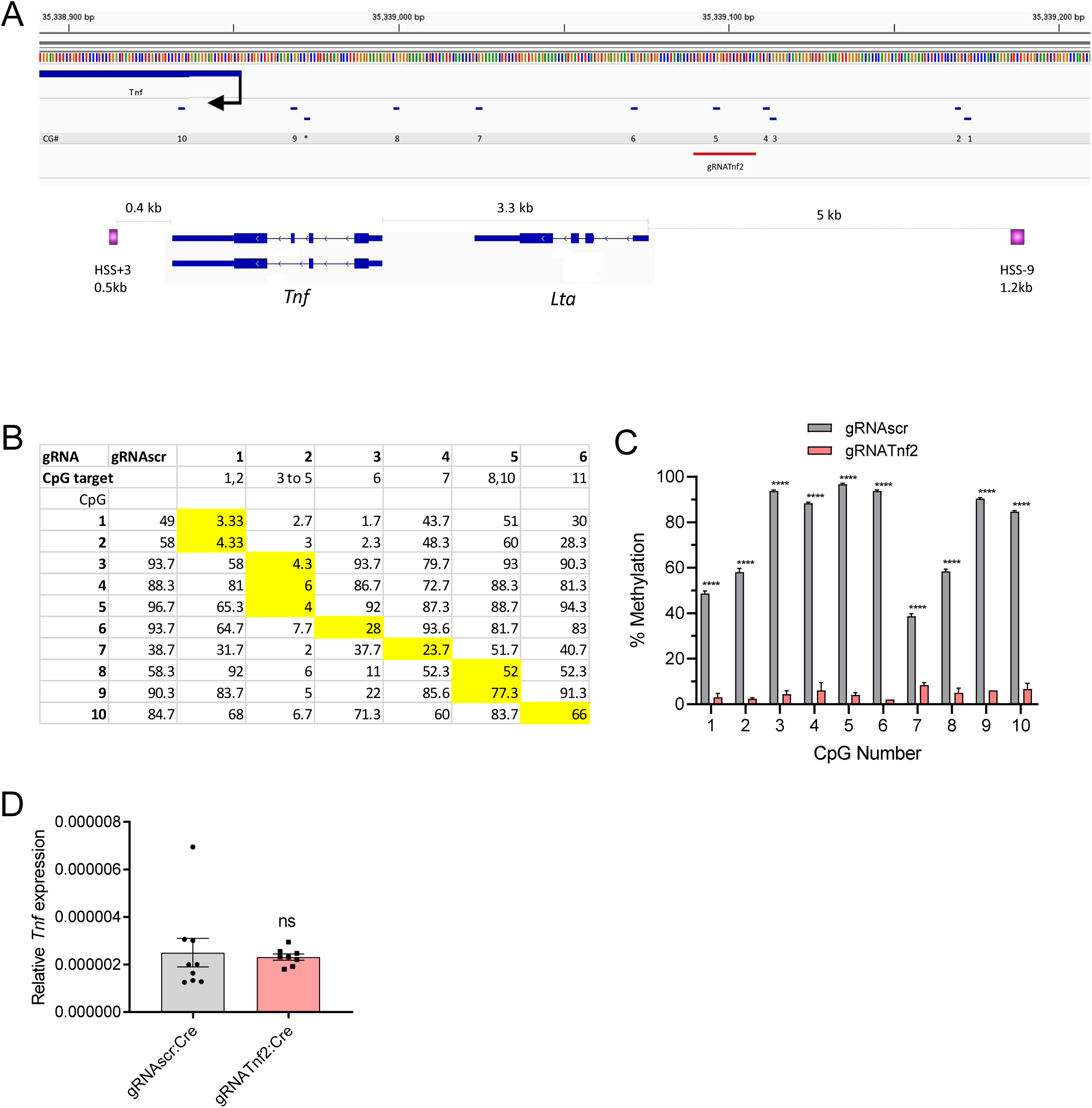
(A) Genome browser view of the murine *Tnf* locus;(Top) each CG location marked by a blue dash and numbered below, TSS indicated by a black arrow., and the location of gRNA*Tnf2* is labeled with a red line and marked accordingly; (Bottom) Two known distal enhancers of *Tnf* expression indicated with purple boxes, named and marked with distances to *Tnf* TSS [67]. (B) Table demonstrating the average methylation of *Tnf* CpGs numbered in (A) as measured by bisulfite-pyrosequencing a function of six candidate *Tnf*-targeting gRNAs or gRNAscr control in NIH-3T3 cells also stably expressing dCas9. CpGs within the gRNA binding site are indicated below the gRNA number and their methylation status is highlighted in yellow in the corresponding gRNAs. (C) DNA methylation levels assessed by bisulfite-pyrosequencing of NIH-3T3 cells stably expressing dCas9 and either gRNA*Tnf2* (pink) or gRNAscr (grey) (n=3). (D) *Tnf* expression in NIH-3T3 cell lines subcloned from those in Fig. 6H stably expressing gRNAscr (grey) or gRNA*Tnf2* (pink) under high-puromycin conditions in combination with dCas9, followed by dCas9 removal by Cre recombinase as assayed by RT-qPCR and normalized to *Actb* expression (n=8-9). * indicates statistically significant difference of p<0.05, ** of p<0.01, *** of p<0.001, **** of p<0.0001, and ns = not significant (Student’s t-test, with Holm-Sidak correction if number of tests is greater than 3).

**Supplementary Table S1.**
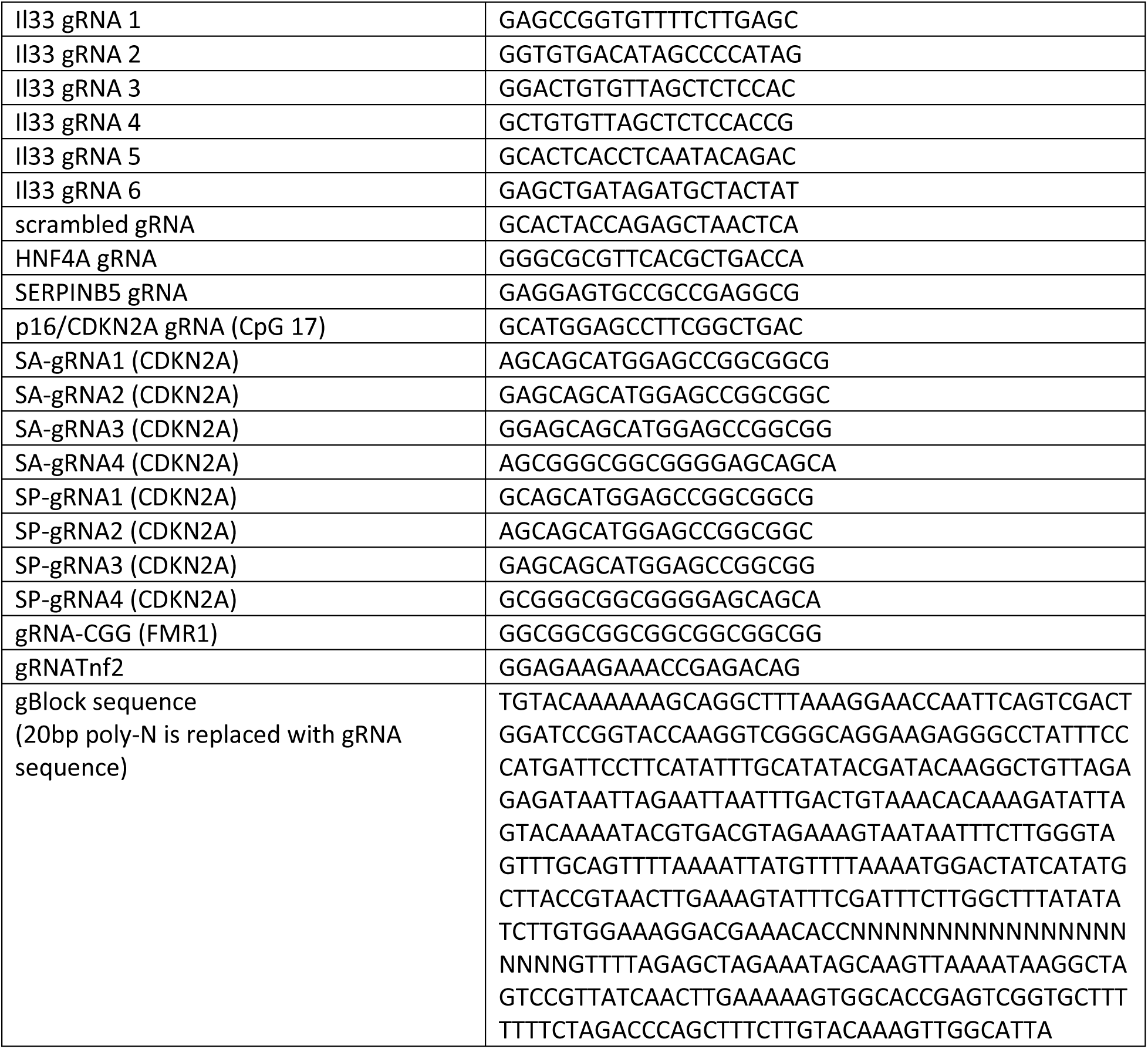
Target sequences of all gRNAs used in the study.

**Supplementary Table S2.**
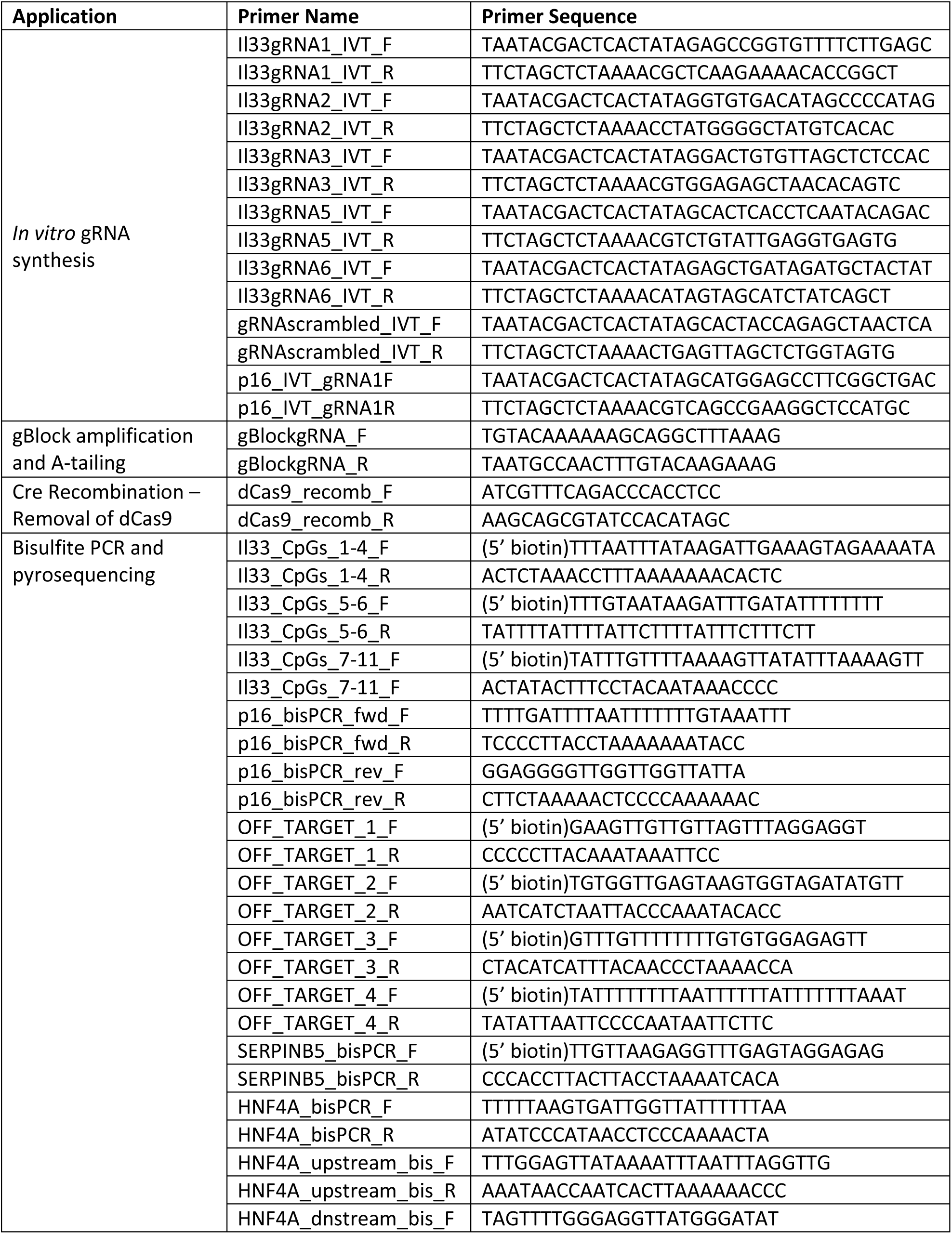
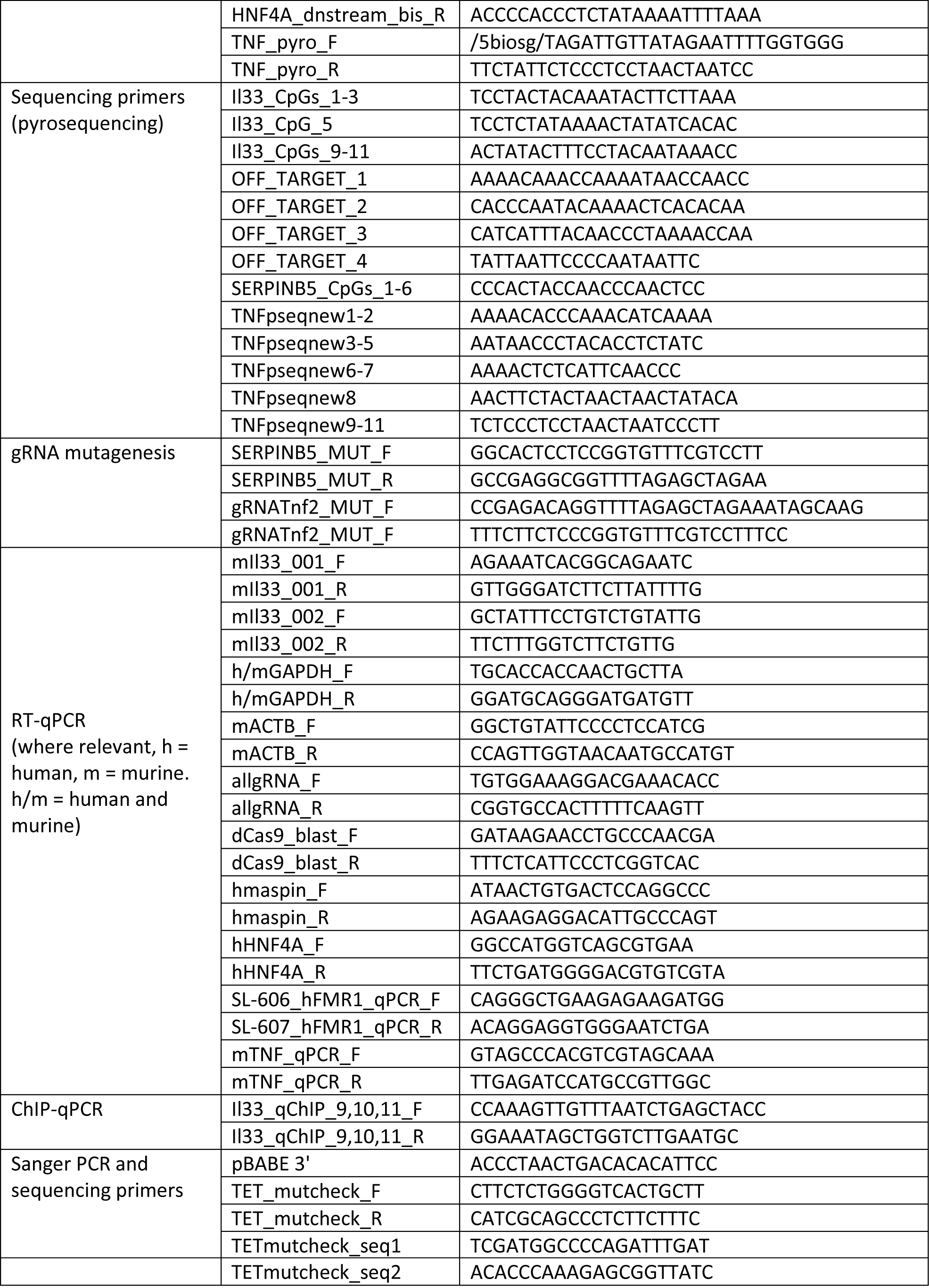
Names and sequences of oligonucleotide primers used in this study.

**Supplementary Table S3.**
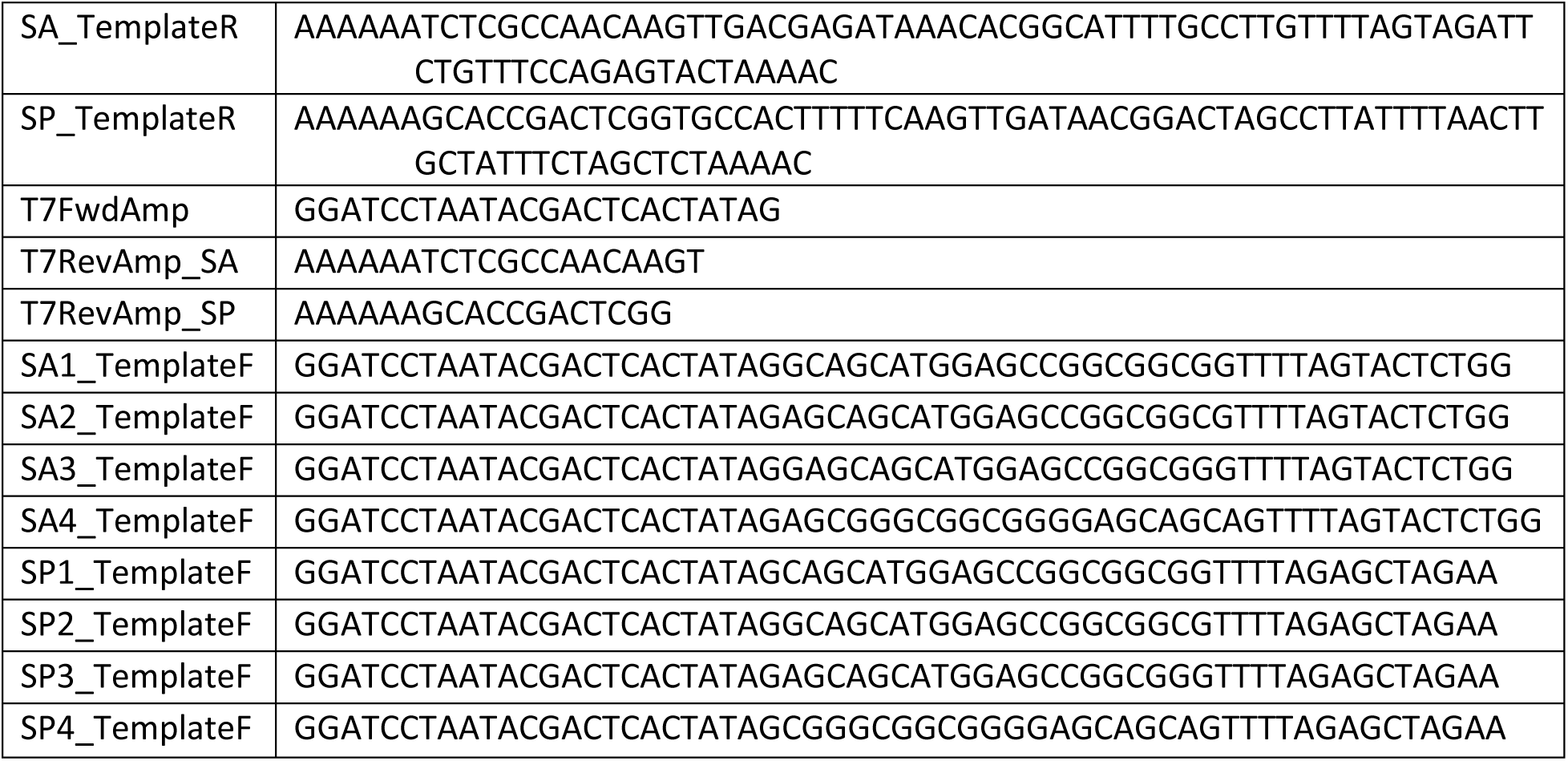
Primers for S. aureus-based strategy for *in vitro* gRNA transcription.

**Supplementary Table S4.**
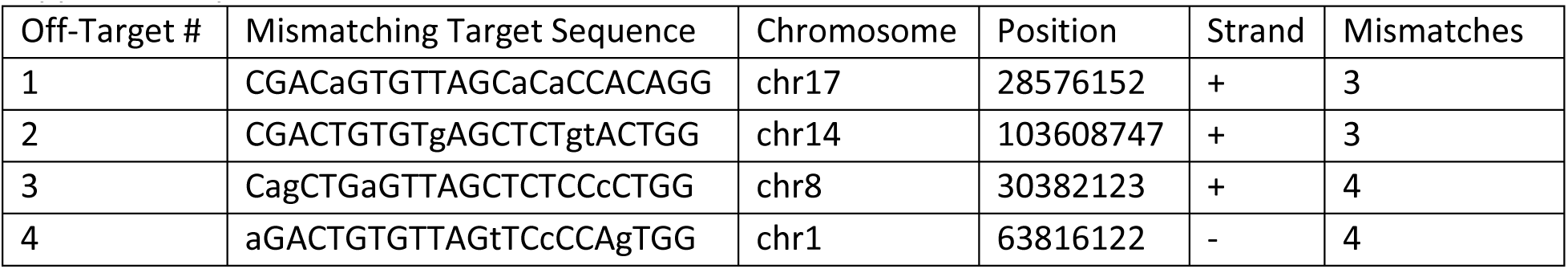
Sequences and locations of predicted mismatched off-target sites for Il33 gRNA3.

**Supplementary Table S5.** Candidate genes in NIH-3T3 cells for robust induction by 5-aza-2’-deoxycytidine. Expression normalized to *ACTB* and expressed as fold change from water-treated controls.

|  |  |  | Normalized to<br>Actb |  | Fold Change |
| --- | --- | --- | --- | --- | --- |
| Gene | 5-aza 1 μM | Control | 5-aza 1 μM | Control | |
| Tm9sf4 | 38.74 | 37.79 | 2.78658E-07 | 2.58E-07 | 1.1 |
| Tnf | 31.34 | 34.205 | 4.70645E-05 | 3.1E-06 | 15.2 |
| Slc44a1 | 24.045 | 22.945 | 0.007391075 | 0.007599 | 1.0 |
| Spata2l | 28.915 | 28.38 | 0.00025275 | 0.000176 | 1.4 |
| Cacna1b | 29.735 | 29.69 | 0.000143168 | 7.08E-05 | 2.0 |
| Fam170a | 40.295 | 40.695 | 9.48353E-08 | 3.45E-08 | 2.8 |
| Nars | 20.15 | 20.16 | 0.109956134 | 0.052374 | 2.1 |
| Atp9a | 36.025 | 39.01 | 1.82965E-06 | 1.11E-07 | 16.5 |
| Mkln1 | 22.535 | 22.175 | 0.021050525 | 0.012958 | 1.6 |
| Cenpw | 24.515 | 23.22 | 0.005336095 | 0.00628 | 0.8 |
| Fam20a | 27.255 | 26.325 | 0.000798732 | 0.00073 | 1.1 |
| Msi2 | 23.55 | 22.595 | 0.010416396 | 0.009685 | 1.1 |
| Taf3 | 24.16 | 23.74 | 0.006824788 | 0.00438 | 1.6 |
| Neb | 39.28 | 37.95 | 1.91653E-07 | 2.31E-07 | 0.8 |
| Foxj1 | 35.96 | 35.91 | 1.91397E-06 | 9.5E-07 | 2.0 |
| Brinp3 | 38.435 | 38.11 | 3.4426E-07 | 2.07E-07 | 1.7 |
| Mcf2 | 32.795 | 32.48 | 1.7167E-05 | 1.02E-05 | 1.7 |
| Il33 | 35.65 | 35.14 | 2.37276E-06 | 1.62E-06 | 1.5 |
| Actb | 16.965 | 15.905 |  |  |  |

